# Cathepsin X is a conserved cell death protein involved in algal response to environmental stress

**DOI:** 10.1101/2024.05.15.594278

**Authors:** Avia Mizrachi, Mai Sadeh, Shifra Ben-Dor, Orly Dym, Chuan Ku, Ester Feldmesser, Amichai Zarfin, John K. Brunson, Andrew E. Allen, Robert E. Jinkerson, Daniella Schatz, Assaf Vardi

## Abstract

Phytoplankton play a crucial role in global primary production and can form vast blooms in aquatic ecosystems. Bloom demise and the rapid turnover of phytoplankton are suggested to involve programmed cell death (PCD) induced by diverse environmental stressors. However, fundamental knowledge of the PCD molecular components in algae and protists in general remains elusive. Previously, we revealed that early oxidation in the chloroplast predicted subsequent cell death or survival in isogenic subpopulations that emerged following H_2_O_2_ treatment in the diatom *Phaeodactylum tricornutum*. Here, we performed transcriptome analysis of sorted sensitive oxidized cells and resilient reduced cells, to discover genes linked to their contrasting fates. By cross-comparison with a large-scale mutant screen in the green alga *Chlamydomonas reinhardtii*, we identified functionally relevant conserved PCD gene candidates, including the cysteine protease cathepsin X/Z (*CPX*). *CPX* mutants in *P. tricornutum CPX1* and *C. reinhardtii CEP12* both exhibited profound resilience to oxidative stress, supporting a conserved function in algal PCD. *P. tricornutum cpx1* mutants, generated using CRISPR-Cas9, also exhibited resilience to the toxic diatom-derived infochemical cyanogen bromide. Phylogenetic and predictive structural analyses show that *CPX* is highly conserved in eukaryotes, and algae of the green and red lineages exhibit strong structural similarity to human cathepsin *CTSZ*. *CPX* is expressed by diverse algae across the oceans and during toxic *Pseudo-nitzschia* blooms, supporting its ecological importance. Elucidating PCD components in algae sheds light on the evolutionary origin of PCD in unicellular organisms, and on the cellular strategies employed by the population to cope with stressful conditions.

## Introduction

Phytoplankton are unicellular photosynthetic organisms that are responsible for ∼50% of global primary production^1^. Yet, they account for less than 1% of the biomass on earth, a gap that is attributed to their high turnover rates^2, 3^. Phytoplankton serve as the base of marine food webs, and are central to the biogeochemical cycling of important elements such as carbon, nitrogen, phosphate, iron and silica^4–8^. They can form massive blooms, which are characterized by “boom and bust” dynamics, with rapid proliferation followed by a synchronized demise in response to diverse environmental stressors^9, 10^. However, we still lack fundamental knowledge regarding how the demise is coordinated.

The induction of programmed cell death (PCD) in response to different environmental stressors has been suggested as an important mechanism contributing to the fast turnover of phytoplankton and rapid bloom demise^11^. PCD is an active and genetically regulated autocatalytic cell death process, as opposed to unregulated necrotic death or death by grazing. In algae, there is accumulating evidence for PCD characteristics in response to various stressors, such as nutrient limitation, heat stress, pathogens, high light, various infochemicals and oxidative stress^12–20^. Reactive oxygen species (ROS) were shown to play an important role in stress sensing and mediating PCD in algae and across the kingdoms of life^21–24^. In the diatom *Phaeodactylum tricornutum*, H_2_O_2_ treatment leads to cell mortality with characteristics of PCD, including externalization of phosphatidylserine, DNA laddering, and compromised cell membranes^16^. Similarily, H_2_O_2_ also induces PCD charactaristics in the green alga *Chlamydomonas reinhardtii*, including membrane permeabilization, alteration of mitochondrial membrane potential, induction of protease activity, and DNA laddering^17^. However, very little is known about the molecular mechanisms involved in algal PCD and in protist PCD in general, as they lack most of the canonical components of well-studied metazoan PCD pathways. Due to this gap of knowledge, the roles and evolutionary origins of PCD in unicellular organisms remain enigmatic.

The evolution of genes with lethal phenotypes, such as PCD-related genes, in unicellular organisms requires that they are tightly regulated and activated only in a subset of cells within the population. In the context of the bloom, during the demise most of the population dies, but some cells are able to survive and serve as the seed for the next bloom. Our current understanding of the mechanisms that mediate acclimation to environmental stressors in marine microorganisms, including algae, is derived primarily from observations at the population level. However, averaging the phenotypes of a whole population can mask the co-existence of distinct subpopulations that employ diverse cellular strategies which can improve the survival of the population as a whole^25–27^. There is accumulating evidence for heterogeneity in algae and specifically in marine diatoms, such as in cell size and morphology^7, 28^, chain length^29^, metabolic states^30–33^, oxidative stress sensitivity^34^, life stages^35–37^, spore formation following viral infection^38^, and more. Studying phenotypic variability within phytoplankton populations can unravel unknown acclimation strategies to environmental stress conditions.

We recently revealed how *P. tricornutum* cells differed in their chloroplast redox state and subsequent cell fate in response to H_2_O_2_, exposing distinct subpopulations of sensitive and resilient cells^34^. Following treatment, a subpopulation of sensitive cells exhibited high oxidation in the chloroplast within minutes, as measured by the redox sensor roGFP^39^ targeted to the chloroplast (chl-roGFP), and eventually died 24 h later. On the other hand, the resilient subpopulation maintained a more reduced state and was able to recover and survive. These subpopulations emerged from isogenic *P. tricornutum* cultures, exposing phenotypic variability in oxidative stress sensitivity. Since chl-roGFP oxidation precedes and predicts cell death in this experimental setup at the single cell level, it can be used to discriminate these subpopulations early on while they are still viable.

Here, we aim to identify the yet unknown PCD genes in algae, and to gain evolutionary insights regarding the origins and conservation of PCD in this important group. We used a unique experimental setup in order to compare transcriptional responses of sensitive and resilient *P. tricornutum* subpopulations to H_2_O_2_, and to identify genes directly linked to their specific fates of survival and cell death. We then implemented a mutant screen in the green alga *C. reinhardtii* as a heterologous system to detect functionally relevant conserved PCD genes for downstream analyses. The cysteine protease cathepsin X/Z (denoted here as CPX) emerged as a PCD related gene. We further characterized its mutants in *P. tricornutum* and *C. reinhardtii*, and studied its phylogeny, structure predictions, and ecological significance in algal blooms.

## Results & Discussion

### *P. tricornutum* sensitive and resilient subpopulations are at distinct transcriptional states following oxidative stress

To directly link between cell fate and gene expression, we performed transcriptome analysis of sorted sensitive and resilient *P. tricornutum* subpopulations. We treated *P. tricornutum* cells expressing chl-roGFP^40^ with 80 µM H_2_O_2_, a dose which leads to induction of cell death in only a fraction of the population after 24 h^34^. Following H_2_O_2_ treatment, two distinct subpopulations were detected: sensitive cells with highly oxidized chl-roGFP and more reduced resilient cells (Fig. S1E-F). These subpopulations were FACS sorted based on chl-roGFP oxidation 2.7 h post treatment, as were control untreated cells, for subsequent transcriptomics analysis (Fig. 1A and Fig. S1). Previously, we demonstrated that cells exhibiting high chl-roGFP oxidation soon after H_2_O_2_ treatment undergo commitment to subsequent cell death^34^. The selected time-point in our current experiment corresponds to a stage in which cells are already committed to either death or survival, though early in the cell death cascade when cells are still viable and do not display PCD characteristics yet^16, 34^. Since the subpopulations emerged from the same genetic background and were exposed to the same conditions but had contrasting fates, the comparison between them can reveal transcriptional patterns that are linked to PCD or stress survival while discriminating them from a general stress response (Fig. 1B).

**Figure 1.**
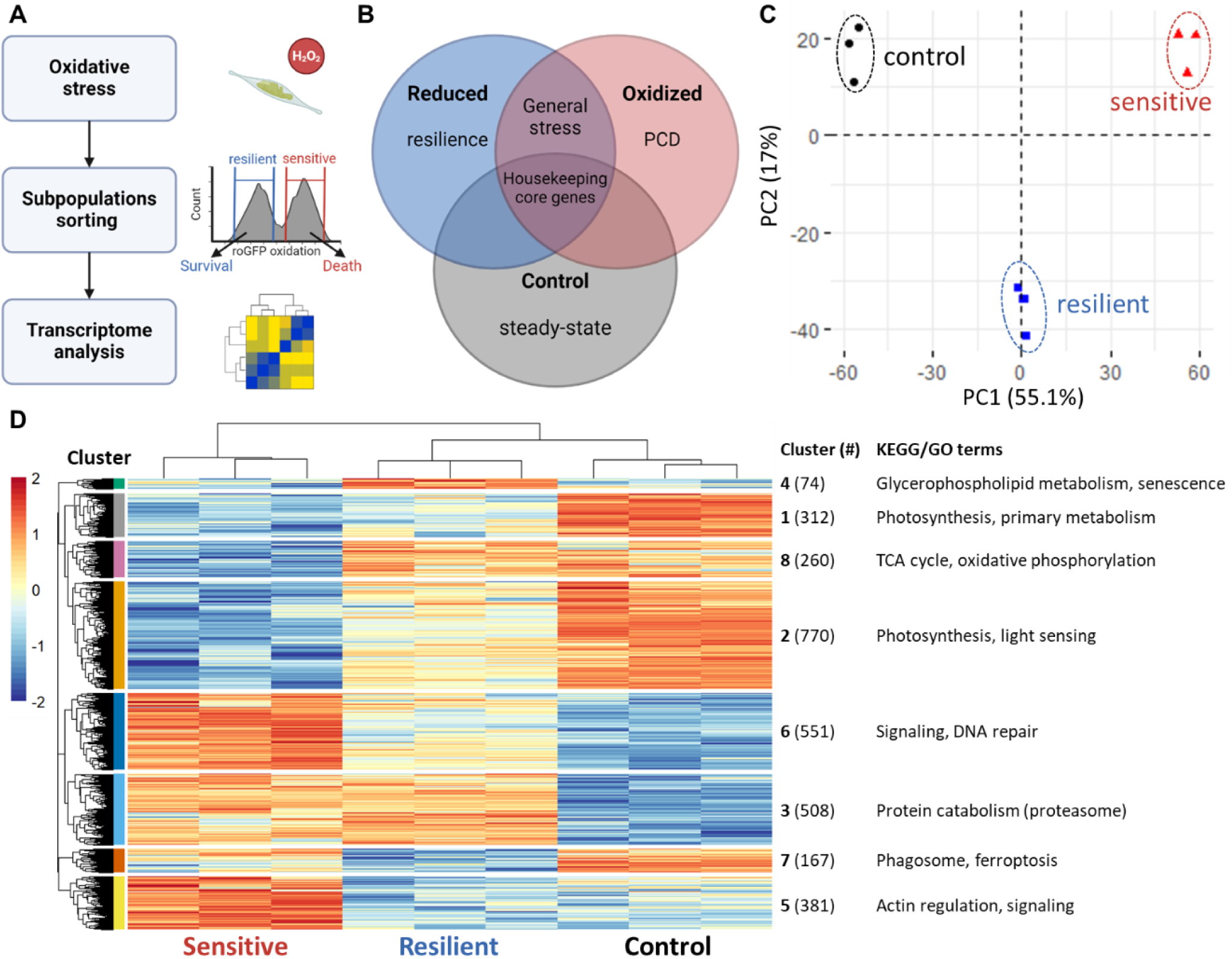
Transcriptome analysis of sorted sensitive and resilient *P. tricornutum* subpopulations following oxidative stress. (***A***) Experimental layout. *P. tricornutum* cells expressing chl-roGFP were treated with 80 µM H2O2, and then sensitive oxidized cells and resilient reduced cells were FACS sorted for transcriptome analysis 2.7 hours later based on chl-roGFP oxidation, as were control untreated cells. (***B***) Conceptual representation of copmparative transcriptomics. As both the sensitive and resilient subpopulations were exposed to the same stress conditions, the comparison between them enables discrimination of gene expression patterns that are linked directly to their specific fates of cell death or stress survival, excluding general stress responses. *(**C**)* The first two components of principal component analysis (PCA) of sorted sensitive (red), resilient (blue) and control (black) samples based on the 6,211 gene transcripts that passed initial filtration for sufficient expression (see Methods). *(**D**)* Gene expression heatmap based on transcriptome analysis of sorted subpopulations, depicting 3,023 differentially expressed genes. Color represents the normalized log2 unique molecular identifier (UMI) count (see Methods), scaled per row. Each row represents a single gene, each column a single sample. Genes and samples were arranged based on Euclidean distances as shown by the dendograms, and genes were clustered into 8 clusters. On the right: cluster number, in parentheses the number of genes within the cluster, and some of the enriched KEGG and GO terms from the analysis in Fig. S3. (*C-D*) The experiment was done with 3 biological repeats of 20,000 sorted cells per sample.

Out of 12,178 coding genes in the *P. tricornutum* genome^41^, 6,211 were detected and 3,023 were significantly differentially expressed in at least one comparison (Fig. 1C-D, Fig. S2 and Table S1, see Methods). Principal components analysis (PCA) demonstrated that biological replicates within each subpopulation clustered together and were clearly separated from the other groups (Fig. 1C). This was also shown by clustering of differentially expressed genes into 8 clusters based on Euclidean distances of their normalized expression across samples (Fig. 1D). The expression patterns specific to each subpopulation clearly demonstrate that each subpopulation is at a distinct transcriptional state, and that both are different from the control.

The clustering revealed genes that exhibited higher or lower expression specific to resilient (clusters 4 and 7 respectively, Fig. 1D) or sensitive cells (clusters 5 and 8 respectively), providing promising leads for identifying genes linked to specific cell fates. To gain functional insights into each cluster, we performed analyses for enriched terms of Gene Ontology (GO) and the Kyoto Encyclopedia of Genes and Genomes (KEGG) (Fig. 1D and Fig. S3). Cluster 5 included genes that had higher expression specifically in the sensitive subpopulation that was committed to cell death, and therefore may be linked to PCD. Among the enriched KEGG and GO terms in this cluster were terms linked to signaling pathways and actin regulation (Fig. 1D and Fig. S3). On the other hand, cluster 8, which included transcripts with lower expression in sensitive cells, was enriched with terms linked to core carbon metabolism (Fig. 1D and Fig. S3), which may represent the shutoff of these processes as part of the cell death cascade. A metabolic shutoff was also detected in the general oxidative stress response, as primary metabolism and photosynthesis terms were enriched in clusters 1 and 2 that exhibited downregulation in both H_2_O_2_-treated subpopulations (Fig 1D and Fig. S3). Known stress-related terms were enriched in clusters that were upregulated in both subpopulations, such as proteasome and protein catabolism in cluster 3 and MAPK signaling and DNA damage repair in cluster 6 (Fig. 1D and Fig. S3). This supports the validity of the data as these clusters are expected to be part of a general oxidative stress response. To summarize, the subpopulations that emerged from genetically homogenous cells, and that were discriminated by their chloroplast redox state, exhibited unique stress-related gene expression signatures. For downstream analyses, we focused on cluster 5 to identify the unknown genes involved in algal PCD.

### Identifying conserved PCD candidate genes using *C. reinhardtii* mutant phenotypes of *P. tricornutum* orthologs

Next, we aimed to identify and characterize conserved PCD genes in distant photosynthetic protists to gain insights into the evolutionary origin of PCD in this important group. We utilized a recent genome-wide mutant library in the green alga *C. reinhardtii*^42, 43^ as a heterologous large-scale functional screen. This library consists of >58,000 indexed insertion mutants that were screened under >120 conditions, including nutrient limitations, exposure to infochemicals, H_2_O_2_, Rose Bengal (which induces singlet oxygen production), grazing and more^42^. The barcoded mutants were pooled together and their relative growth was measured under different conditions to characterize their phenotypes.

We hypothesized that inactivation of a gene involved in PCD may result in increased resilience to some stress conditions, as was shown in yeast^44–47^. Therefore, we selected *P. tricornutum* genes that were specifically induced in the sensitive subpopulation that was committed to cell death (cluster 5), and that the *C. reinhardtii* mutants of their orthologs exhibited a resilient phenotype, meaning that they grew better than the rest of the population under specific treatments (Fig. 2, Table S2). Such resilient phenotypes in mutants are considered rare and can provide strong supportive evidence for a pro-PCD function. We also focused on resilient mutants specifically under oxidative stress, since it may show a more direct link between *C. reinhardtii* and *P. tricornutum* in this setup. This approach enabled us to link gene expression and phenotypes using a heterologous system and to identify functionally relevant candidates for downstream analyses.

**Figure 2.**
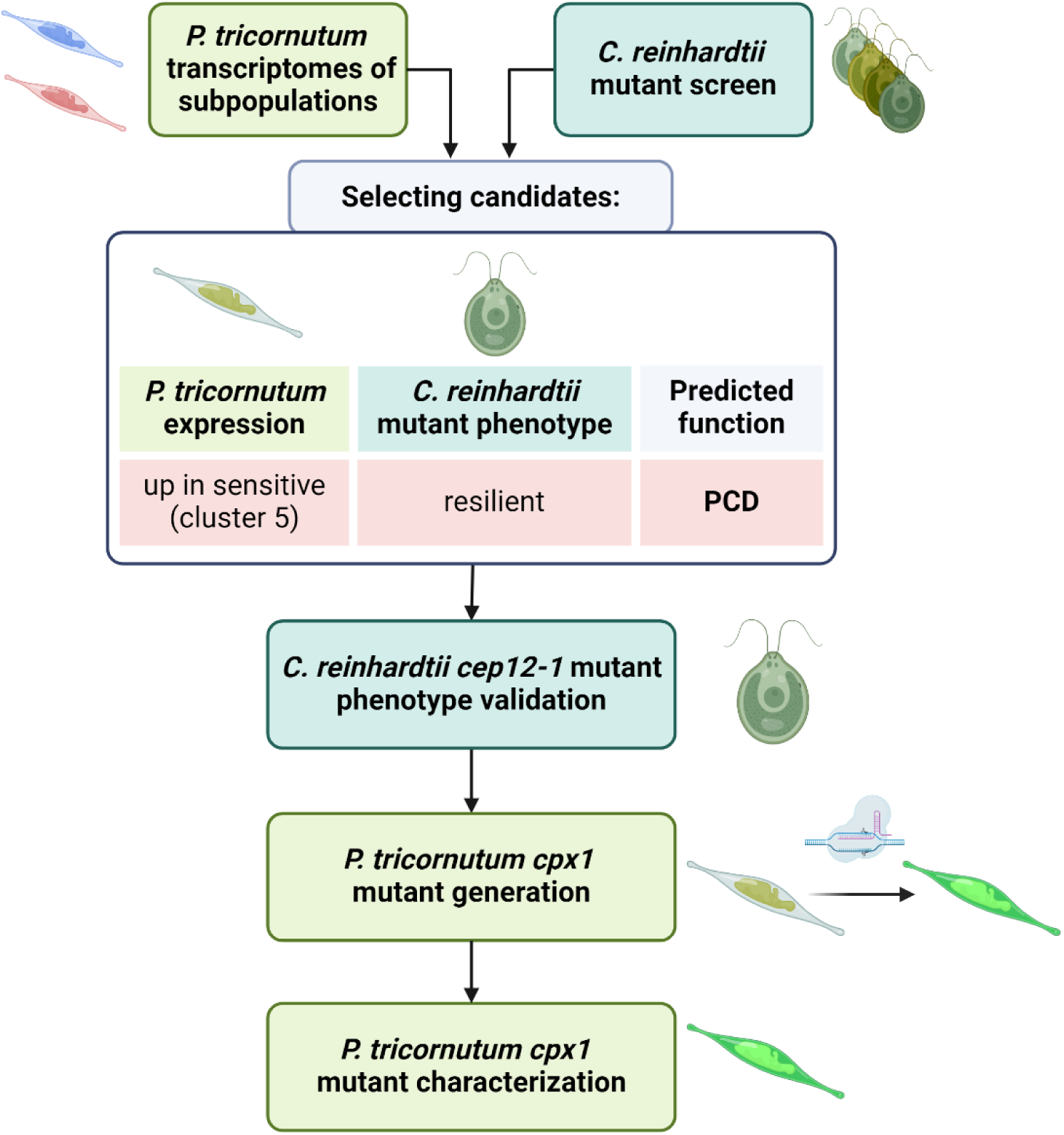
Schematic representation of identification and characterization of PCD candidates in *P. tricornutum* and *C. reinhardtii*. To identify conserved candidate genes involved in PCD in algae, we combined expression data from *P. tricornutum* sorted subpopulations (Fig. 1) and a published large-scale mutant screen performed in *C. reinhardtii*^42^. We selected genes that were 1) induced specifically in sensitive *P. tricornutum* cells that were subsequently going to die, and thus suspected to be involved in PCD (cluster 5 in Fig. 1D), and 2) *C. reinhardtii* mutants of their orthologs exhibited a resilient phenotype in at least one condition in the screen, preferably related to oxidative stress (Table S2). *CPX* emerged as a PCD related gene, and was selected for further investigation both in *C. reinhardtii cep12-1* mutant and in *P. tricornutum cpx1* mutants that were generated using CRISPR-Cas9.

The comparative analysis revealed 21 hits (Table S2), among them the papain-type cysteine carboxypeptidase *CPX* – CYSTEINE ENDOPEPTIDASE 12 (*CEP12*, encoded by Cre12.g498850) in *C. reinhardtii* and *CPX1* (Phatr3_J9664) in *P. tricornutum*. The *C. reinhardtii* mutant *cep12-1* was found in the top <0.5% mutants when comparing their relative growth following H_2_O_2_ treatment to control conditions (Table S2). Cathepsins are a conserved protein family found across eukaryotes, and they are involved in protein cleavage in a wide variety of pathways such as proteolytic activity within lysosomes, signaling pathways, immunity, and in different types of PCD in plants and animals^48–52^. Among the family of cathepsins, CPX was studied mainly in mammalian systems in the context of various diseases^53–55^, while its function and roles in algae and in other protists remain completely unknown. Since other cathepsin types were shown to be involved in PCD in plants and animals^48–52^, we chose to study its role in the response to oxidative stress and PCD induction in *C. reinhardtii* and *P. tricornutum*.

### *C. reinhardtii cep12-1* mutant is resilient to H_2_O_2_

To validate its phenotype and to further characterize *C. reinhardtii cep12-1* mutant, we investigated its response to H_2_O_2_ treatment. Cells were treated with lethal doses of H_2_O_2_ that were shown to induce PCD characteristics in *C. reinhardtii*^17^, and cell death was measured over time by Sytox Green staining, which selectively penetrates dying cells, using flow cytometry (Fig. 3A and Fig. S4A). *C. reinhardtii cep12-1* mutant had a significantly lower fraction of dead cells 24 h after treatment with 1.75-3 mM H_2_O_2_ (Fig. 3A) and reached its maximal percentage of dead cells at a later time point compared to the WT (Fig. S4A). To further investigate the physiological response of *C. reinhardtii cep12-1* mutant to oxidative stress, we examined its maximal photosystem II (PSII) efficiency (Fv/Fm) following H_2_O_2_ treatment (Fig. S4B). Under control conditions, Fv/Fm values of *cep12-1* mutant were similar to the WT, demonstrating that PSII is not compromised in the mutant under steady state conditions. In response to H_2_O_2_ treatment, the decrease in Fv/Fm values was more profound in the WT, while *cep12-1* maintained higher values (Fig. S4B). Overall, these results demonstrated that *C. reinhardtii cep12-1* mutant exhibits resilience to H_2_O_2_, supporting the potential role of CEP12 in PCD following oxidative stress.

**Figure 3.**
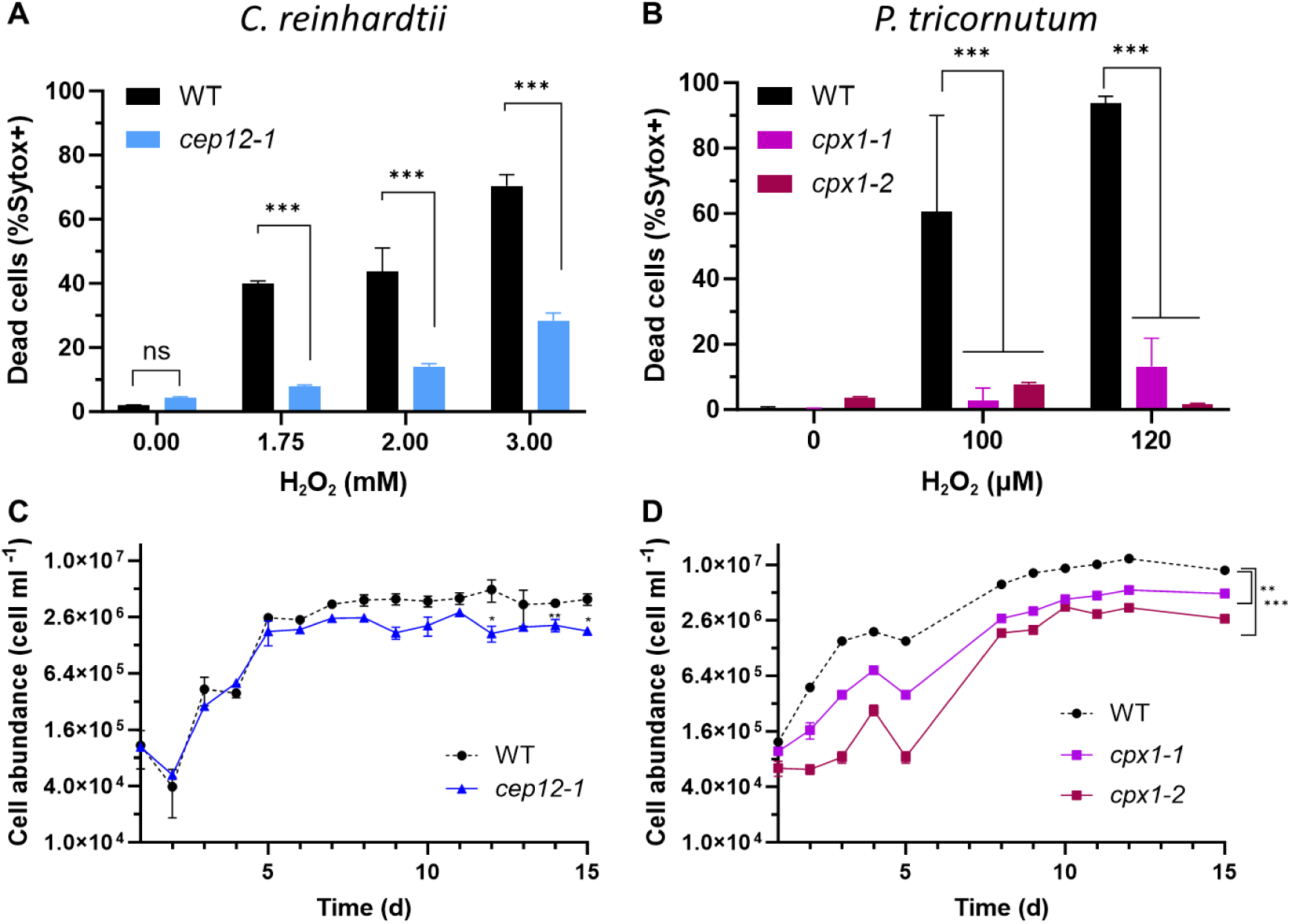
*CPX* mutants in both *C. reinhardtii* and *P. tricornutum* are resilient to oxidative stress. Mortality in response to H_2_O_2_ treatments (***A-B***) and growth under control conditions (***C-D***) of *C. reinhardtii cep12-1* mutant (A, C; blue) and *P. tricornutum cpx1* mutants (B, D; *cpx1-1* – purple, *cpx1-2* – red) compared to their corresponding WT strain (black). (***A-B***) The percentage of dead cells 24 h post H_2_O_2_ treatment as measured by Sytox Green staining using flow cytometry. (***A-D***) Values represent the mean ± sd, N=3. Statistical significance was calculated by using Two-way ANOVA (A-B) and repeated measures ANOVA (C-D) as compared to WT. Asterisks represent *P*-values: *<0.05, **<0.01, ***<0.001.

### Resilience to oxidative stress is conserved in *P. tricornutum cpx1* mutants

Next, we sought to link the results obtained in *C. reinhardtii* to *P. tricornutum* by generating *CPX1* deletion mutants in *P. tricornutum*. Two independent mutants were generated employing CRISPR-Cas9 with two guide RNAs (gRNAs) targeting different regions in the *P. tricornutum CPX1* gene in order to induce a large deletion that is likely to disrupt its function (see Methods, Fig. S5 and S6). In *cpx1-1*, one allele had a deletion of 879 base pairs (bp) that included the active site, while the other allele had an insertion of 2 bp leading to a frame shift and a premature stop codon before the catalytic site (Fig. S5 and S6). In *cpx1-2*, both alleles had 711 bp deletion that included the active site (Fig. S5 and S6). Based on the sequence, *CPX1* is expected to be non-functional in both mutant lines. In terms of growth, both *C. reinhardtii cep12-1* and *P. tricornutum cpx1* mutants exhibited lower maximal carrying capacity at stationary phase (Fig. 3C and D respectively), while *cpx1* mutants also had lower cell abundance at most time points, suggesting CPX has an additional role during growth.

Next, we examined the response of *P. tricornutum cpx1* mutants to oxidative stress. *P. tricornutum cpx1* and WT cells were treated with different doses of H_2_O_2_, and cell death was quantified 24 h later using flow cytometry as described for *C. reinhardtii.* Since *P. tricornutum* is much more sensitive to H_2_O_2_ than *C. reinhardtii*, we used lower H_2_O_2_ doses that are similar to those used in previous works^16, 17, 34^. The fraction of dead cells was significantly lower in response to all examined concentrations of H_2_O_2_ in both *cpx1* mutants compared to the WT (Fig. 3B), in accordance with the resilience to H_2_O_2_ of *C. reinhardtii cep12-1* (Fig. 3A).

In addition, we examined the response of *P. tricornutum cpx1* mutants to environmentally-relevant concentrations of cyanogen bromide (BrCN), a diatom-derived infochemical that induces oxidative-stress mediated cell death^16, 56^. In *P. tricornutum*, BrCN induces early oxidation of roGFP targeted to the chloroplast, nucleus and mitochondria, which is followed by cell death^16^. In response to 4 µM BrCN, both *cpx1* mutants had a significantly smaller fraction of dead cells compared to the WT (Fig. S7). When treated with 5 µM BrCN, *cpx1-2* had significantly lower mortality while *cpx1-1* was not significantly different from the WT (Fig. S7). Thus, *cpx1* mutants exhibited increased resilience towards BrCN. This suggests that CPX1 is involved in oxidative stress-related PCD also during biotic interactions that are mediated by toxic infochemicals such as BrCN.

### CPX is highly conserved from algae to humans

To gain a broader perspective on the evolution and conservation of CPX, we generated a phylogenetic tree of the cathepsin domain using representative species from different lineages that were within the top BLAST hits of *P. tricornutum* CPX1 (Fig. 4A, Table S3, see Methods). The analysis resulted in five major cathepsin groups: three subgroups of CPX (denoted here CPX-I, CPX-II and CPX-III), cathepsin B, and a more distant group of other cathepsins which can be viewed as an outgroup (Fig. 4A). CPX subgroups had a mixed phylogenetic distribution within them. Multiple species had more than a single CPX gene, sometimes distributed between different subgroups. For example, *C. reinhardtii* CEP12 was in the CPX-III subgroup, closer to metazoan CPX than to *P. tricornutum* CPX1 or to another *C. reinhardtii* CPX protein (CEP6, XP_001702410.2) that was found in CPX-I (Fig. 4A). Furthermore, some genes had two CPX domains, each of them in a different clade (denoted as “a” and “b” in Fig. 4A, Table S4). On the other hand, most metazoans formed a clade within the CPX-III group, without other CPX subtypes. Notably, all diatoms in our analysis had at least two CPX genes, and all formed a highly conserved clade within the CPX-II group, including *P. tricornutum* CPX1. We detected 4 CPX genes in *P. tricornutum* (Fig. 4A), but only CPX1 was induced specifically in cells that were committed to undergo PCD (cluster 5 in Fig. 1D, see Table S5), suggesting they have different functions. These patterns could be explained by multiple gene duplication and diversification events, as was suggested in other cathepsin types^57, 58^, leading to the expansion of the CPX gene family.

**Figure 4.**
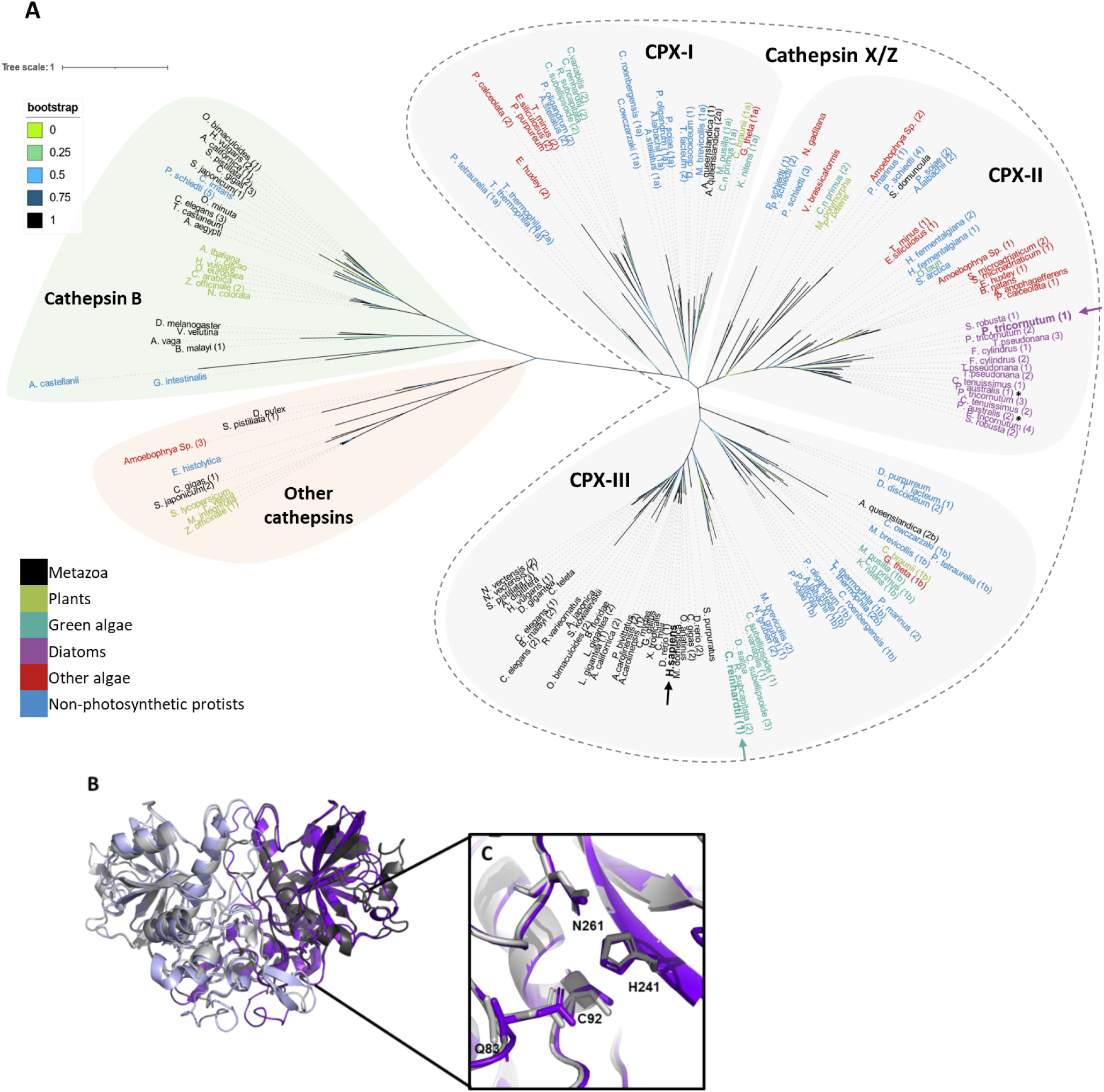
Conservation of CPX across the tree of life. **(*A*)** Unrooted phylogenetic tree of the conserved cathepsin domain of homologs of *P. tricornutum* CPX1. Branch color represents bootstrap values. Background shade represents the cathepsin group: CPX subgroups CPX-I, CPX-II, and CPX-III (grey), cathepsin B (green), and others (orange). Label colors represent organism group: black – metazoans; green – plants; cyan – green algae; purple – diatoms; red – other algae; blue – non-photosynthetic protists. Arrows pointing at *P. tricornutum* CPX1 (purple), *C. reinhardtii* CEP12 (cyan) and *Homo sapiens* (black) are shown. * *P. australis* (1) and (2) represent contigs from the metatrascriptome from Brunson *et al.*^60^ shown in Fig. 5C. (***B-C***) Overlays of AlphaFold2 predicted model of *P. tricornutum* CPX1 dimer (purple) and the experimental structure of the human CTSZ dimer (PDB ID 1EF7, grey). The monomers in each homodimer are shown in light and dark shades. (***C***) Zoom in on the conserved active site residues of human CTSZ and *P. tricornutum* CPX1 that are shown in stick grey and purple respectively. The numbering of the human CTSZ active site residues: Q83, C92, H241 and N261 are labeled, corresponding to *P. tricornutum* CPX1 residues: Q79, C88, H259 and N281 respectively.

Interestingly, we did not detect CPX in vascular plants, most arthropods, fungi, bacteria and archaea. CPX was likely lost in vascular plants, in which the closest homologs were found in the cathepsin B group, while it still exists in liverwort and moss as well as in green algae (Fig. 4A). This is in contrast to the expansion and diversification that occurred in other cathepsins in vascular plants, mainly cathepsin L-like proteases^58, 59^. This conservation pattern supports that CPX evolved early in the evolution of the first eukaryotes, and was likely already present in the last eukaryotic common ancestor (LECA) as was recently suggested by another phylogenetic analysis of cathepsins^57^. Later, it was likely lost in several groups, including fungi, vascular plants and in most Arthropods.

To investigate CPX protein structural conservation, we examined the similarity between the predicted protein structures of *P. tricornutum* CPX1 and *C. reinhardtii* CEP12, comparing them to the most similar published CPX structure of *Homo sapiens* cathepsin X/Z (CTSZ, PDB code 1EF7)^61, 62^. Human CTSZ is biosynthesized as an inactive zymogen, and requires *in trans* activation by another protease such as cathepsin L^63^. The active mature protein acts as a homodimer with monopeptidyl carboxypeptidase activity^61, 63, 64^, unlike most other cathepsins, which usually act as monomers and possess endopeptidase activity. We generated structural models of CPX dimers using AlphaFold2 employing sequences corresponding to the experimentally determined structure of human CTSZ (see Methods). The top-ranking predicted models of *P. tricornutum* CPX1 and *C. reinhardtii* CEP12 revealed high agreement with the experimental dimer structure of human CTSZ, with RMSD values of 0.593 Å and 0.519 Å respectively (Fig. 4B and Fig. S8), demonstrating their high structure similarity. *C. reinhardtii* CEP12 exhibited greater similarity to human CTSZ than to *P. tricornutum* CPX1, evidenced by its higher sequence identity (Fig. 4A and Fig. S9) and closer resemblance in dimer structures (Fig. 4B and Fig. S8). This is also demonstrated by the conservation pattern of cysteine residues that are predicted to be involved in disulfide bond formation (marked with green numbers in Fig. S9). Moreover, many residues in human CTSZ involved in the homodimer interface (highlighted by boxes in Fig. S9) are strictly conserved across all three proteins. While certain residues are not conserved among the three proteins, they are still predicted to play a role in dimer formation (Fig. S9). The active site residues, Q83 and the catalytic triad C92, H241 and N261 in Human CTSZ, demonstrated strict conservation in *P. tricornutum* CPX1 (Q79, C88, H259 and N281, respectively, Fig. 4D) and in *C. reinhardtii* CEP12 (Q93, C102, H250 and N272, respectively), as indicated by blue arrows in Fig. S9. Protein targeting also seems to be conserved, as human CTSZ is targeted to lysosomes but can also act extracellularly^65^, and *P. tricornutum* CPX1 was also predicted to be localized to the lysosome, along with almost half of the CPX proteins used in the phylogenetic tree analysis (Table S3). *C. reinhardtii* CEP12 was predicted to be localized to the extracellular space, with lysosome targeting as the second best hit. Taken together, the sequence and overall predicted structural similarity suggest a shared biochemical activity among these proteins, and demonstrate high conservation across eukaryotes of distant lineages.

### CPX is widely expressed by diverse algae in the marine environment and during a toxic diatom bloom

In order to assess the environmental significance of CPX, we examined its expression in the marine ecosystem. First, we investigated the expression of homologs of the conserved domain of *P. tricornutum* CPX1 in TARA oceans metatranscriptome data using the Ocean Gene Atlas^66^ (Fig. 5A-B). CPX was expressed in diverse sampled locations and by a wide variety of marine protists, including stramenopiles (13% of the reads, mainly diatoms), dinoflagellates (39%) and haptophytes (8%, Fig. 5A-B). Next, we investigated the expression of CPX1 homologs in a recent time-series study of *Pseudo-nitzschia australis* dominated toxic blooms in Monterey Bay, California^60^. These blooms were sampled in high temporal resolution with near-weekly sampling during 2015, for metatranscriptome analysis coupled with analysis of domoic acid (DA) toxin production. We detected two *P. australis* annotated contigs, PaCPX1 and PaCPX2, which had high homology to *P. tricornutum* CPX1, with ≥65% protein similarity (E-value ≤3e-112, Fig. S10). These contigs also clustered together with *P. tricornutum* CPX1 in the phylogenetic tree (marked with asterisks in Fig. 4A). There were two major bloom events during the examined period of April 15, 2015 to September 30, 2015. Both CPX contigs were detected throughout that time, and had different expression dynamics (Fig. 5C), suggesting CPX has an important function also in natural diatom blooms. PaCPX1 expression profile increased concomitant with *Pseudo-nitzschia* cell abundance, peaking prior to demise events. During the second bloom event, the peaks of PaCPX1 and *Pseudo-nitzschia* cell abundance were also accompanied by high DA concentrations (Fig. 5C). Notably, PaCPX1 reaches an expression peak on July 1^st^, coinciding with substantial silicate and iron exhaustion in Monterey Bay^60^. PaCPX1 clustered together with other genes linked to stress in module 5 in the WGCNA analysis performed by Brunson *et al.*, characterized by genes with annotations of protein turnover, chaperones, antioxidants and in addition cathepsin A^60^. PaCPX2 had a lower relative expression in the first bloom event, and had a more pronounced increase in the summer bloom event, during which it also peaked together with cell counts prior to the crash. The high conservation of CPX, its wide expression by a large variety of algae in the world’s oceans, and its expression patterns during a toxic *P. australis* bloom suggest that CPX plays a role in diverse natural algae populations.

**Figure 5.**
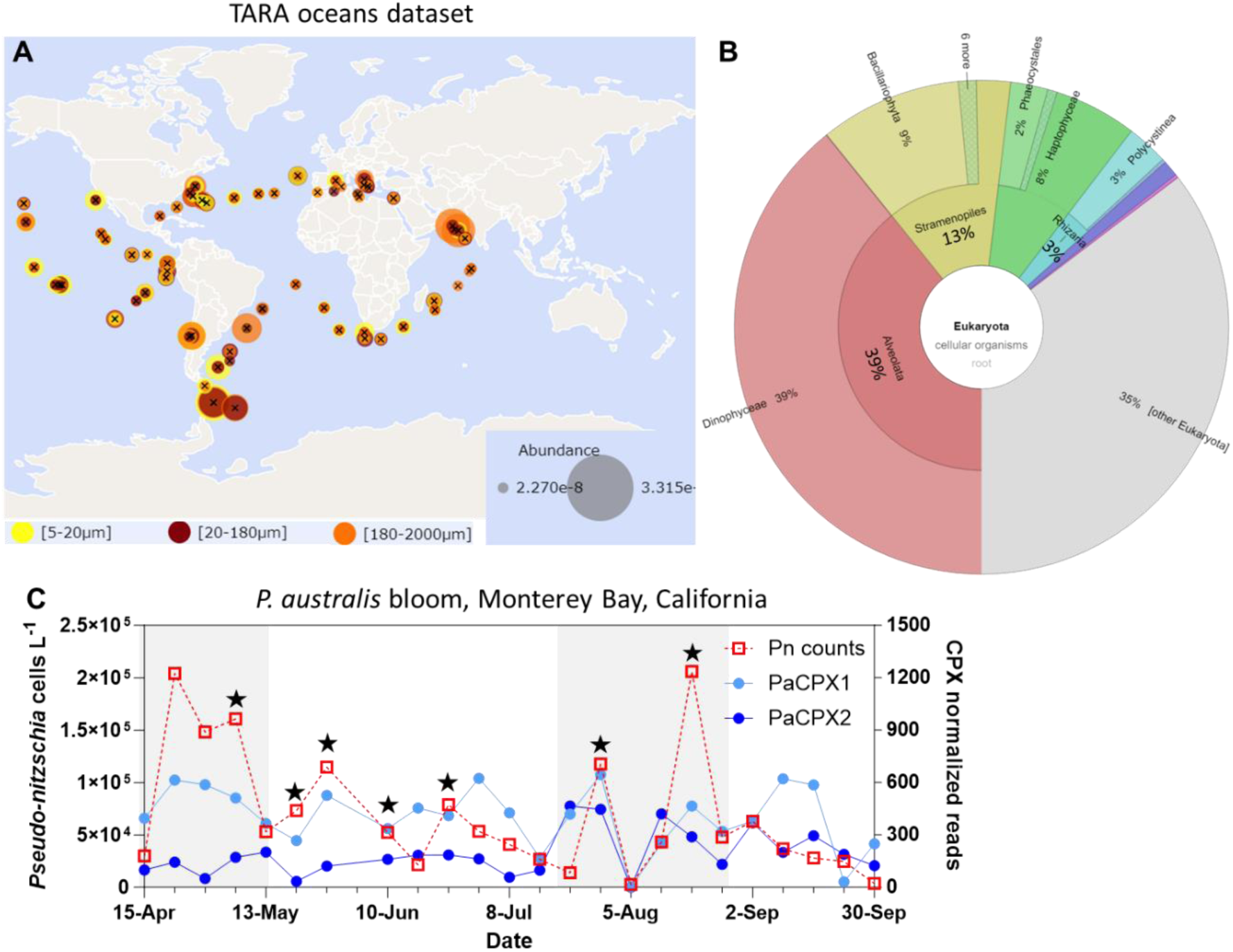
Expression of CPX in natural marine environments. (***A-B***) Expression of contigs with high homology to *P. tricornutum* CPX1 conserved domain (E-value < 1E-40) in TARA Oceans eukaryotic metatranscriptomic data analyzed using the Ocean Gene Atlas^66^. (***A***) Relative expression of CPX1 homologs in surface seawater across the world oceans. Circle size represents transcript abundance normalized as percent of total reads. Color represents the size fraction. (***B***) Krona visualization of taxonomic distribution of CPX1 eukaryotic homologs in TARA Oceans metatranscriptomics data. Red – Alveolata (dinoflagellates, 39%); yellow – Stramenopiles (13%, mainly diatoms); green – Haptophyceae (8%); cyan – Rhizaria (3%); grey – other eukaryotes (35%). (***C***) Normalized expression of *P. australis* CPX1 homologs (blue and light blue) and *Pseudo-nitzschia* cell counts (Pn counts, red) during the bloom period (April 15, 2015 to September 30, 2015) in Monterey Bay, California, data from Brunson *et al.*^60^. Grey background marks the two major bloom events. Black stars mark days with particulate domoic acid >900 ng/L. Raw reads were normalized to total *P. australis* read counts, multiplied by 1 x 10^6^.

### A conserved role for CPX in cell fate regulation

The phylogenetic analysis and structural predictions of CPX demonstrate high conservation between very distant phylogenetic groups of metazoans and photosynthetic protists from the green and red lineages. This conservation pattern together with the shared resilient phenotypes of *C. reinhardtii cep12-1* and *P. tricornutum cpx1* mutants suggest that CPX may have an important role in cell fate regulation in a wide variety of protists of distant lineages and potentially also in metazoans. Interestingly, CPX of algae from both the green and red lineages are related to animal CPX, while vascular plants are missing this gene. Recent studies in mammalian systems demonstrated various phenotypes of CPX (CTSZ) mutants^67^, including increased resilience to certain drugs^68, 69^ and pathogens^70, 71^. Animal CTSZ is involved in immunity, cellular senescence, cancer and other pathologies^65, 72–74^. Though it was suggested to be involved in various PCD pathways in humans^75^, it was not directly demonstrated, and its role in animal PCD remains elusive. Cathepsins of other types were shown to be involved in PCD in animals and plants^48–52^, and here we demonstrate the involvement of CPX in algal PCD. Interestingly, the induction of different cathepsins, including CPX, correlated with the induction of cell death in a dinoflagellate during co-culture with an algicidal bacterium^76^. A potential PCD-related activity of CPX could be, for example, through cleavage and activation of proteins involved in the PCD signaling cascade following its release from damaged lysosomes, as shown for cathepsin B^49, 52^. To summarize, we propose that the role of cathepsins in PCD also evolved early in the evolution of eukaryotes, and that cathepsins can be used to study the evolution of PCD machinery from unicellular to multicellular organisms.

### The origin of PCD in unicellular organisms

How unicellular organisms evolved lethal traits such as PCD remains an open question. In multicellular organisms, PCD has important functions in development, maintaining homeostasis, and immunity. Protists lack most of the canonical PCD machinery known from metazoans, such as caspases, p53, and Bcl2, and we lack fundamental knowledge of any of their PCD molecular components. Nevertheless, PCD characteristics have been demonstrated in algae, other protists and bacteria^13–16, 18, 19, 77, 78^. Here, we reveal the first PCD gene in aquatic protists with conserved function between distant algae from the red and green lineages.

The emergence of PCD in diverse unicellular organisms raises fundamental evolutionary questions regarding its function on the cellular and population levels. Several hypotheses were proposed to explain how PCD genes could have evolved in unicellular organisms, for example to prevent the expansion of viral infection, or when PCD in a subpopulation can help the survival of the entire population^15, 77, 79, 80^. Such mechanisms can provide a supply of nutrients under starvation conditions, and may be favorable when there is strong kin selection^81^. For example, supernatant of *C. reinhardtii* cells that were exposed to heat stress that led to PCD enhanced the population growth more than medium derived from non-PCD lysed cells^15^. In order to coordinate PCD at the population level, ROS can be used to communicate stress signals between cells^21^. PCD serves an important role in multicellular structures or during cooperative behaviors, and it was suggested as a key step in the evolution of multicellularity^82, 83^. The observed cell-to-cell heterogeneity in the physiological response of diatoms to environmental stressors can provide an acclimation strategy by the induction of PCD in a subpopulation, mediated by CPX, while allowing the survival of the remaining cells of the same population.

Another explanation for the evolution of PCD genes in unicellular organisms could be that they have a dual role. While under certain conditions they have vital functions, at different circumstances they lead to a lethal phenotype. For example, yeast Mca1 (also known as Yca1) was shown to have dual biochemical activity, as it switches between proteolytic activity that promotes cell death and chaperone-like activity during replicative aging through a mechanism that involves binding of calmodulin^84, 85^. CPX may also have a dual function, as suggested by the impaired carrying capacity at stationary phase in CPX mutants in both *C. reinhardtii* and *P. tricornutum*, and by the lower cell abundance at most time points measured in *P. tricornutum cpx1* mutants (Fig. 3C-D). This result may indicate a tradeoff between oxidative stress resilience and cell growth, and suggests that CPX has an additional important pro-survival role. CPX may function, for example, in protein turnover and the recycling of their building blocks during stationary growth phase or under nutrient starvation, while having a lethal function promoting PCD under strong oxidative stress.

### The ecological importance of algal PCD

The ecological consequences of PCD and its effects on nutrient recycling in microbial food webs await further investigation^86^. PCD has the potential to make cellular contents more accessible to some groups rather than others^15^, for example by providing nutrients to neighboring conspecifics, or serving as carbon sources for bacteria in the microbial loop instead of being grazed upon by zooplankton. In addition, PCD characteristics and stress were associated with increased transparent exopolymeric particles production^12, 87–90^. Thus, it may affect aggregation and sinking rates, which in turn may affect the bioavailability of cellular biomass and carbon sequestration in the deep ocean. Genes linked to PCD have the potential to be used in future studies as biomarkers, detecting early stages of phytoplankton cell death prior to bloom demise.

Furthermore, cell death is associated with the release of cellular contents to the seawater, including toxins and other infochemicals^91^ that can act as signals of stress surveillance systems during bloom succession^92^. Thus, PCD genes may be used together with other molecular biomarkers, such as toxin biosynthesis genes as a proxy to detect early toxin release, as was recently demonstrated^60^. The identification and characterization of PCD genes could be used to improve our understanding of cellular strategies employed during bloom dynamics and to study the downstream consequences of PCD in an ecological and biogeochemical context^11^. In the future, the approach taken in this study could also be used to identify other conserved genes that are involved in stress resilience, oxidative stress sensing and more.

In conclusion, this work provides novel insights on the unknown molecular machinery of PCD in photosynthetic protists and its ecological significance during algal bloom succession in the ocean.

## Methods

### Culture maintenance

*P. tricornutum* culture, accession Pt1 8.6 (CCMP2561 in the Provasoli-Guillard National Center for Culture of Marine Phytoplankton) was purchased from the National Center of Marine Algae and Microbiota (NCMA, formerly known as CCMP). Cultures were grown in filtered sea water (FSW) supplemented with F/2 media (trace metals, Fe citrate, vitamins, NaH_2_PO_4_·2H2O and NaNO3^93^) at 18°C with 16:8 hours light:dark cycle and light intensity of 170 μmol photons m^-2^ sec^-1^ supplied by cool-white LED lights (Edison, New Taipei, Taiwan). Unless otherwise specified, all experiments were performed with exponentially growing cultures (∼5·10^5^-1·10^6^ cell ml^-1^ on the day of the experiment). For measuring growth rates, cultures were diluted to 1·10^5^ cells ml^-1^ on day 0 of the experiment. *C. reinhardtii cpx12-1* strain (Cre12.g498850; LMJ.RY0402.222468) from the CLiP library and CC325 which was used as the WT strain in this study were purchased from the Chlamydomonas Resource Center. Cassette insertion in *cpx12-1* was validated using Polymerase Chain Reaction (PCR) according to the instructions of the Chlamydomonas Resource Center. *C. reinhardtii* cultures were grown in liquid TAP medium^94^, under light intensity of 40–60 µmol photons m^-2^ sec^-1^ at 25°C.

### Sorting of *P. tricornutum* subpopulations for transcriptome analysis

*P. tricornutum* cells expressing chl-roGFP^40^ were treated with 80 µM H_2_O_2_ and sensitive oxidized cells and resilient more reduced cells were then FACS sorted 2.7 h later using BD FACS AriaII based on chl-roGFP oxidation. Gates used for sorting, population hierarchy and representative FACS plots are shown in Fig. S1. Untreated control cells were sorted immediately after H_2_O_2_ treated samples based on positive roGFP signal regardless of their oxidation state (the parent gate of “oxidized” and “reduced” subpopulations).

Chlorophyll autofluorescence to detect chlorophyll-containing cells was measured using 488 nm excitation (ex) and 695/40 emission (em). Relative roGFP oxidation was measured using the ratio between two fluorescence channels, i405 (ex 405 nm, em 530/30) and i488 (ex 488 nm, em 530/30), as the roGFP ratio (i405/i488) increases upon oxidation of the probe^95^. As references for maximal roGFP oxidation and reduction, treatments of 200 µM H_2_O_2_ and 2 mM Dithiothreitol were used respectively. To keep the sorting as short as possible, 20,000 cells were sorted per sample. Samples were sorted directly into cold lysis buffer, in Eppendorf™ LoBind tubes that were kept on ice prior to sorting and during sorting at 4°C, and immediately after sorting underwent vortex, spin down, and flash frozen using liquid N_2_. Samples were kept at -80°C until further processing.

### Library preparation and sequencing

cDNA library preparation was done using the MARS-seq protocol as described in^96^, adapted for small amount bulk RNA samples. RNA extraction was conducted using the NucleoSpin® RNA XS kit (Machery-Nagel, Düren, Germany). Then, reverse transcription was done to generate barcoded cDNA containing the unique molecular identifier (UMI) sequences (unique to each mRNA molecule) and the sample ID barcodes (Table S6). Unused primers were then degraded with Exonuclease I (NEB). The cDNA from different samples was then combined into pools, and underwent second-strand synthesis and *in vitro* transcription to amplify the sequences linearly. RNA products were fragmented and ligated with single-stranded DNA adapters containing pool-specific barcodes. A second round of RT was carried out, followed by polymerase chain reaction with primers containing Illumina adapters to construct DNA libraries for paired-end sequencing on a NextSeq machine (Illumina).

### Transcriptome analysis

MARS-seq data was processed using the User-friendly Transcriptome Analysis Pipeline (UTAP)^97^. Reads were trimmed using cutadapt^98^ and mapped to *P. tricornutum* genome annotation version 3 (Phatr3) on EnsemblProtist, assembly ASM15095v2, using STAR^99^ v2.4.2a with default parameters. Reads were demultiplexed based on the pool and sample ID barcodes. The pipeline quantifies the genes annotated in RefSeq (expanded with 1000 bases toward 5’ edge and 100 bases toward 3’ bases). Counting was done using htseq-count^100^. Normalization of the counts and differential expression analysis was performed using DESeq2 R package^101^ with the parameters: betaPrior = True, cooksCutoff = FALSE, and independent Filtering = FALSE. Raw *P* values were adjusted for multiple testing using the procedure of Benjamini and Hochberg. Genes were initially filtered for sufficient expression levels: ≥5 UMI raw counts in at least 3 samples and ≥10 UMI counts in at least one sample. Each UMI may have multiple reads and represents a single transcript copy in the original sample. Out of 12,178 coding genes in the *P. tricornutum* genome^41^, 8214 were detected, 6,211 passed data filtration based on minimum expression levels (see Table S1), and 3,023 were considered as significantly differentially expressed in at least one comparison (is.DE = “yes” in Table S1) according to the following criteria: adjusted *P* value ≤ 0.05 and Log2 fold change ≥1 and baseMean (mean normalized UMI counts across all samples) ≥ 5. GO^102^ and KEGG pathway orthologous groups (KO)^103^ were annotated based on the eggNOG 4.5 databse ^104^ using the eggNOG-mapper ^105^. GO enrichment analysis was done using topGO package in R. GO and KEGG enrichment analyses were done using a hypergeometric test, terms with less than 2 genes were filtered out. Subcellular localization predictions were done using HECTAR^106^ and AsaFind^107^. Analyses were performed in R unless stated otherwise.

### Detecting conserved candidate PCD genes in *C. reinhardtii* and *P. tricornutum*

Orthologs between *P. tricornutum* and *C. reinhardtii* were detected using the protein BLAST best hits that also passed the thresholds of E-value < 1e-3 and bit score ≥ 30. Then, we selected genes that were in cluster 5 (Fig. 1D) in the transcriptome analysis of sorted *P. tricornutum* subpopulations, and that had a resilient phenotype in at least one condition in the *C. reinhardtii* large-scale mutant screen^42^ (see Table S2). A resilient phenotype was considered if the mutant had better relative growth in that condition compared to the other pulled mutants, according to the analysis performed in^42^. In addition, to focus on oxidative stress responses we also included the top 500 mutants (out of >58,000 mutants) under treatments of Rose Bengal and H_2_O_2_.

### Enumeration of algal cell abundance and cell death measurements

All cultures were counted and diluted a day before the experiment and counted again on the day of the experiment to ensure the same cell concentrations between samples. Cell concentration was measured using Multisizer 4 COULTER COUNTER (Beckman Coulter, Indianapolis, USA), except for *P. tricornutum* growth curve in which cells were quantified using the CytoFLEX S flow cytometer (Beckman coulter, Indianapolis, USA). For flow cytometry analysis, the cells population was identified by plotting the chlorophyll fluorescence (excitation: 488 nm, emission: 663 - 737 nm) versus forward scattered (FSC) light (a proxy for cell size). Cell death analysis was done using Sytox Green (Invitrogen), a nucleic acid stain that penetrates compromised membranes, which is a characteristic of dead or dying cells. The Sytox green stock solution was diluted to 5 mM in dimethyl sulfoxide (DMSO) and then added to the samples to a final concentration of 1 μM. Then, samples were incubated in the dark for 30 minutes at room temperature and analyzed by flow cytometry using the green channel (ex: 488 nm, em: 525 nm).

### H_2_O_2_ and BrCN treatments

*P. tricornutum* cells were treated with H_2_O_2_ in 24 well plates (2 ml culture per well), and monitored for cell death as described above. H_2_O_2_ (diluted in DDW) was added to the cultures in final concentrations of 80-120 μM in *P. tricornutum*, and 1-3 mM in *C. reinhardtii*, a concentration range that was used in previous studies and that was shown to induce PCD characteristics in those organisms^16, 17, 34^. BrCN treatment was induced with three doses (1.75, 2.5, 5 μM), and control cells were treated with 1% acetone. After treatment, cells were incubated for 24 hours under normal growth conditions and analyzed for cell death as described above.

### Quantification of photosynthetic efficiency

Photosynthetic efficiency for *C. reinhardtii* cells was measured using the imaging PAM system (Heinz Walz, Effeltrich, Germany) in 24 well plates, 2 ml per well after 5 minutes of dark incubation. Photosynthetic efficiency was determined as Fv/Fm, calculated as previously described^108^. Fm represents the maximum fluorescence emission level in the dark measured with a saturating pulse of light (emission peak at 450 nm, 2,700 μmol photons m^-2^ sec^-1^, 800 ms); Fv = Fm – F0.

### Generation of *P. tricornutum* mutants using CRISPR/Cas9

For transformation of *P. tricornutum* cells, we utilized the CRISPR-Cas9 method^109^. *P. tricornutum cpx1* mutants were created using a set of two guides targeting the *CPX1* gene, transformed into WT cells. Guides were chosen to maximize on-target cleavage and minimize off-target risk using the following programs: MIT guide design^110^ and sgRNAdesigner^111^ in their Benchling implemetations (www.benchling.com), CRISPOR^112^, PhytoCRISP-Ex^113^, and CasOFFinder^114^. CPX1 guide sequences: gRNA1 – GTGTTCTAGTCGAGTGTGAC; gRNA2 – GGACCAATTCACAAATACCC. Transformation was done using a biolistic particle delivery system (Bio-Rad, Hercules, CA, USA) as previously described^115^. Cells were co-transformed with 3 plasmids: 2 plasmids containing diatom-adapted Cas9 and the gRNA^109^ targeting different regions of the CPX1 gene, and a plasmid containing a zeocin resistance cassette used for mutant selection (Fig. S11). Cells were plated on agar plates 2 days prior to bombardment at a concentration of 2x10^7^ cell per plate (50% FSW+F/2, 1.5% agar). M17 W-particles (Bio-Rad, Hercules, CA, USA) were coated with 5 μg DNA in the presence of CaCl_2_ (2.5 M) and spermidine (0.1 M). Cells were bombarded using a rupture disc of 1300 psi, using the hepta-adaptor according to the manufacturer’s instructions. Following the biolistic shooting, cells were transferred to recover on selection plates (50% FSW+F/2, 1.5% agar with 100 μg/ml Phleomycin). Later, colonies that grew on selection plates were isolated and sub-cloned on plates with 100 μg/ml Zeocin (InvivoGen). In order to verify the CRISPR-Cas9 knockout of *CPX1*, cell lysates of antibiotic resistant colonies were prepared in lysis buffer (1% TritonX-100, 20 mM Tris– HCl pH 8, 2 mM EDTA) in an Eppendorf tube by repeated freezing and thawing. Then, cell lysate (5 μl) was used for PCR amplification of the genomic targets with primers specific to the gene of interest. The PCR products were first analyzed by agarose gel electrophoresis. To validate that the mutagenesis caused frameshift insertions or deletions, random colonies were chosen and subjected to PCR amplification and Sanger sequencing of the target region (Fig. S5 and S6). Analysis was done using the sequence alignment programs Sequencher**®** (version 5.4.6 DNA sequence analysis software, Gene Codes Corporation, Ann Arbor, MI USA http://www.genecodes.com) and SnapGene® software (from Dotmatics; available at snapgene.com). Primers used in this study were designed with Primer3^116^, and are detailed in Table S7.

### Cathepsin sequence alignment and phylogenetic tree

Sequences from representative organisms of different kingdoms were chosen from the first 5000 protein Blast^117^ hits using *P. tricornutum* CPX1 as the query. Sequences were then cut to the Peptidase_C1 superfamily domain (cl23744 in CDD^118^). Multiple alignment was performed with Muscle 3.8.31^119^ and a phylogenetic tree was constructed with PhyML (v. 3.0)^120^. The PhyML trees were visualized with iTol (v. 6)^121^.

### Cathepsin X/Z structure models

Multiple sequence alignment was performed using MultAlin^122^. AlphaFold2 algorithm^123, 124^, developed by DeepMind, was employed to generate structural models of *P. tricornutum* CPX1 and *C. reinhardtii* CEP12 dimers. At first, pairwise sequence alignment of the full length (FL) sequences of the different pairs were determined. FL Human CTSZ shares 31.5% sequence identity with *P. tricornutum* CPX1 and 44.1% with *C. reinhardtii* CEP12, while *P. tricornutum* CPX1 shares 27.2% sequence identity with *C. reinhardtii* CEP12 (Fig. S9). Notably, higher pairwise sequence identity was observed when incorporating the sequence used for the experimentally determined structure of Human CTSZ (PDB code 1EF7), and aligning it with *C. reinhardtii* CEP12 and *P. tricornutum* CPX1, excluding the N-terminal and C-terminal regions predicted to be flexible and disorderly by the AlphaFold2 algorithm. The five structural models generated by AlphaFold2 for the FL *C. reinhardtii* CEP12 and *P. tricornutum* CPX1 exhibited flexibility and disorder in both their N-terminal regions (M1-D64 and M1-E57 respectively) and C-terminal regions (G310-Q370 and A311-QF356 respectively), as indicated by their notably low local Distance Difference Test (pLDDT) scores (30%). Therefore, in our model predictions, we utilized *P. tricornutum* CPX1 and *C. reinhardtii* CEP12 sequences corresponding to the experimentally determined structure of Human CTSZ dimer: A65-V311 for *C. reinhardtii* CEP12, and L58-N312 for *P. tricornutum* CPX1 (Fig. S9). In these regions, CTSZ shares 37.6% aa sequence identity with CPX1 and 57.4% with CEP12, while CPX1 shares 37.1% with CEP12 (Fig S9). With these sequences, all five models generated by AlphaFold2 displayed high confidence scores, as reflected by elevated pLDDT values (84-86%), with no significant differences observed among the group of five models for *P. tricornutum* CPX1 and *C. reinhardtii* CEP12. Protein localization was predicted using DeepLoc2^125^, using the high quality setting.

### Geographic distribution of CPX expression in TARA Ocean dataset

To assess the bio-geographic distribution expression of CPX in the natural environment we used Ocean Gene Atlas platform^66^. For the TARA query, we used the MATOUv1+g DATASET with a threshold of 1e-40 and the abundance shown as percent of total reads. The *P. tricornutum* CPX1 domain input was the following sequence: LPMAFSWGNVNGRSYLTKSLNQHIPQYCGSCWAHAALSVLGDRIMIAQSQEEDSSILDEFNLSVQFLLNCAGEYAGSCYGGSTTGVFDFIQDMGYIPYETCQPYLACSDDSDEGICSFVNTTCSPEAICRTCSPDGICQAVTTFPNATVAEYGRYRYELFATMAEIYLRGPVTASIDAGPIHKYPGGVLWDNPKYHSDKTNHAVSIVGWGYDYDEEKQYWIVRNSWGQYWGEMGFFRIELGKNLLKIESNIAWANP.

## Data availability

The MARS-seq dataset and accompanying information have been deposited in NCBI’s Gene Expression Omnibus^126^ and will be made available upon publication through GEO Series accession number GSE266642 (https://www.ncbi.nlm.nih.gov/geo/query/acc.cgi?acc=GSE266642).

## Acknowledgments

We thank Angela Falciatore and Marianne Jaubert for kindly providing us plasmids used for cloning. We thank Shiri Graff van Creveld and Shilo Rosenwasser for valuable comments on the manuscript. This research was supported by the Israeli Science Foundation (ISF) (grant #1972/20) awarded to AV and DS.

## Author Contributions

A.M, M.S, A.V designed the research, analyzed the data, and wrote the article with contributions of the co-authors. A.M conducted the MARS-seq experiment of *P*. *tricornutum*, and performed the analysis with the help of C.K and E.F. R.E.J provided the *C. reinhardtii* mutant screen data. M.S and A.M preformed experiments involving *C. reinhardtii*. A.M and S.B.D designed *P. tricornutum* mutant generation. D.S, M.S, A.Z and A.M carried out the *P. tricornutum* mutant transformation and validation steps, S.B.D and M.S performed mutant sequence analysis. M.S performed experiments with *P. tricornutum* mutants. S.B.D performed the phylogenetic tree analysis, M.S and A.M designed the tree. O.D performed the protein structure prediction analysis. J.K.B and A.E.A provided transcriptomic data of *Pseudo-nitzschia* blooms in Monterey Bay and assisted with data analysis. All authors approved the manuscript.

## Materials & Correspondence

Correspondence and requests for materials should be addressed to A.V.

