## Supplementary information for "Cathepsin X is a conserved cell death protein involved in algal response to environmental stress"

### Supplementary figures

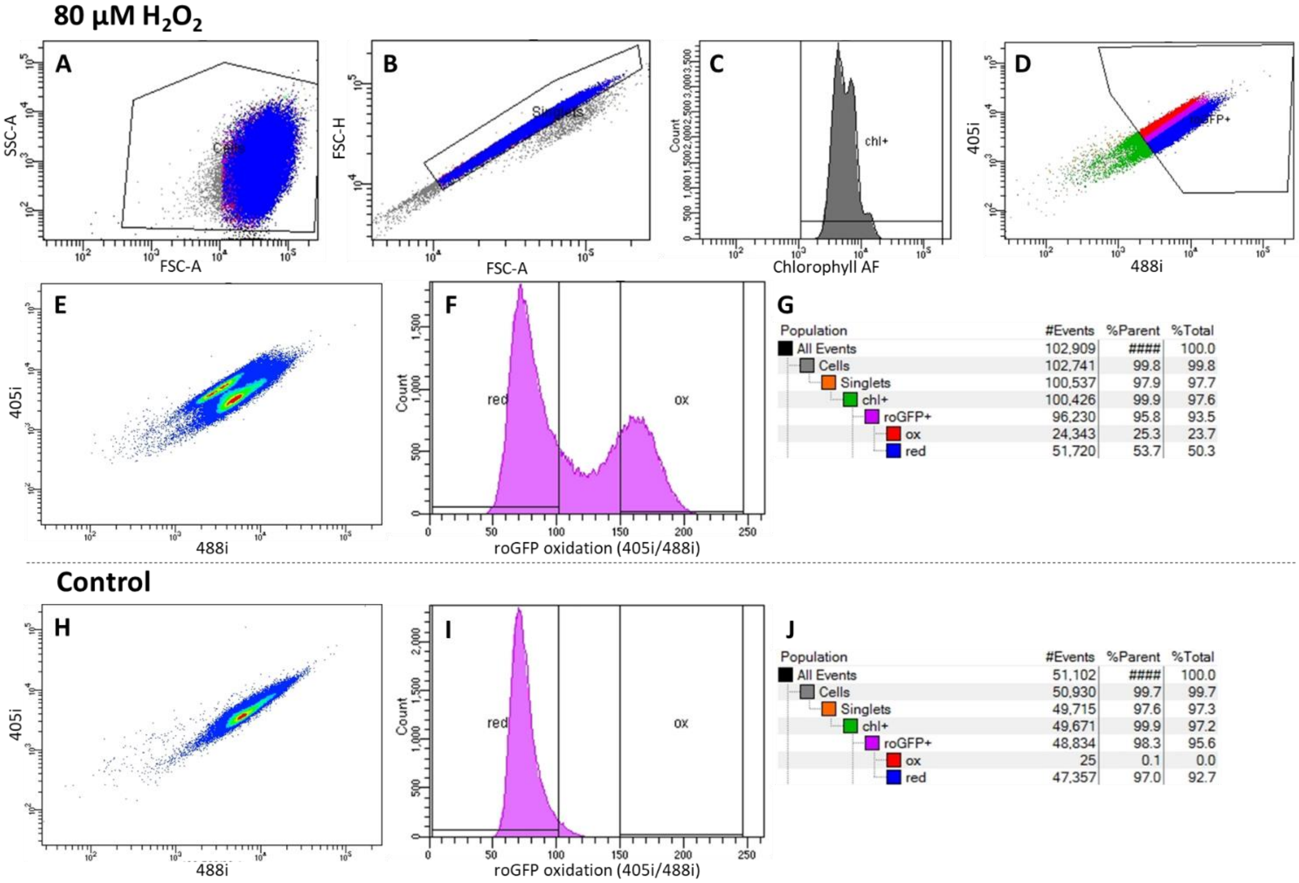


**Figure S1. FACS sorting of *P. tricornutum* subpopulations based on chl-roGFP oxidation.** FACS plots and the gates used for sorting of *P. tricornutum* cells expressing chl-roGFP following treatment with 80 µM $H_2O_2$ (A-G) and in untreated control (H-J). (***A***) Scatterplot of side scatter (SSC) area vs. forward scatter (FSC) area, representing a proxy for size, "cells" gate is shown. (***B***) Scatterplot of FSC height vs. FSC area to exclude doublets, "singlets" gate is shown. (***C***) Chlorophyll auto-fluorescence (AF), "chl+" gate for chlorophyll containing cells is shown. (***D, E, H***) Scatterplot (D) and density plots (E, H) of 405i vs. 488i, representing the roGFP oxidation state. The roGFP ratio (i405/i488) increases upon oxidation of the probe[1]. "roGFP+" gate in D depicts cells with strong chl-roGFP signal regardless of their oxidation state. (***F, I***) The roGFP ratio, higher values represent a more oxidized state. Gates used for sorting sensitive and resilient subpopulations in 80 µM $H_2O_2$ treated samples are shown: "ox" – highly oxidized sensitive cells, "red" – reduced resilient cells. (***G, J***) Population hirerchy, depicting the populations in the different gates with their number of events (#Events) and frequencies out of the parent gate (%Parent) and out of the total events (%Total). In control samples, cells were sorted based on the roGFP+ gate (D) regardless of their oxidation state, but almost all cells exhibited a reduced state (H-J).

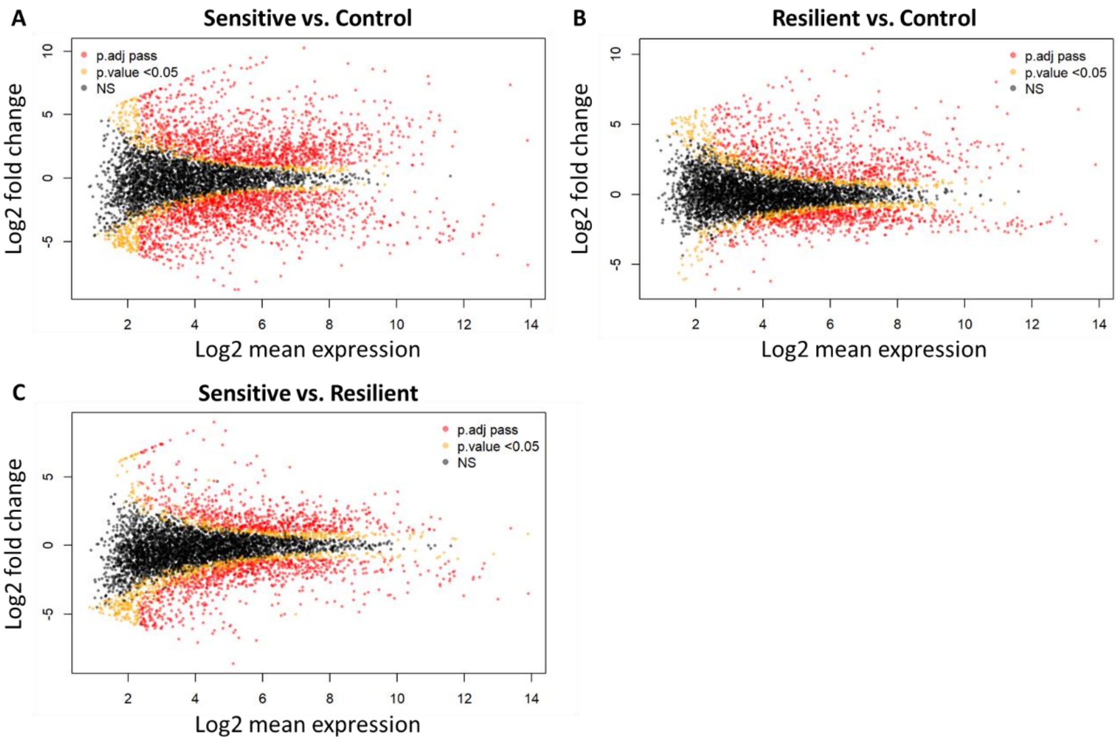


**Figure S2. MA plots of gene expression of sorted *P. tricornutum* subpopulations.** Comparisons of gene expression of *P. tricornutum* sensitive oxidized cells and resilient reduced cells that were FACS sorted based on chl-roGFP oxidation 2.7 h post 80 µM $H_2O_2$ treatment and of sorted untreated control cells. MA plots of the Log2 fold change of normalized UMI counts against the log2 baseMean (mean normalized UMI count across all samples) expression in each comparison: sensitive vs. control (***A***), resilient vs. control (***B***), and sensitive vs. resilient cells (***C***). Red – Differentially expressed genes in the specific comparison that meet the criteria: adjusted *P*-value ≤ 0.05 and Log2 fold change ≥1 and baseMean ≥ 5. Orange – *P*-value < 0.05. Black – non significant. Only genes passing initial data filtration for sufficient expression levels are shown (see Methods).

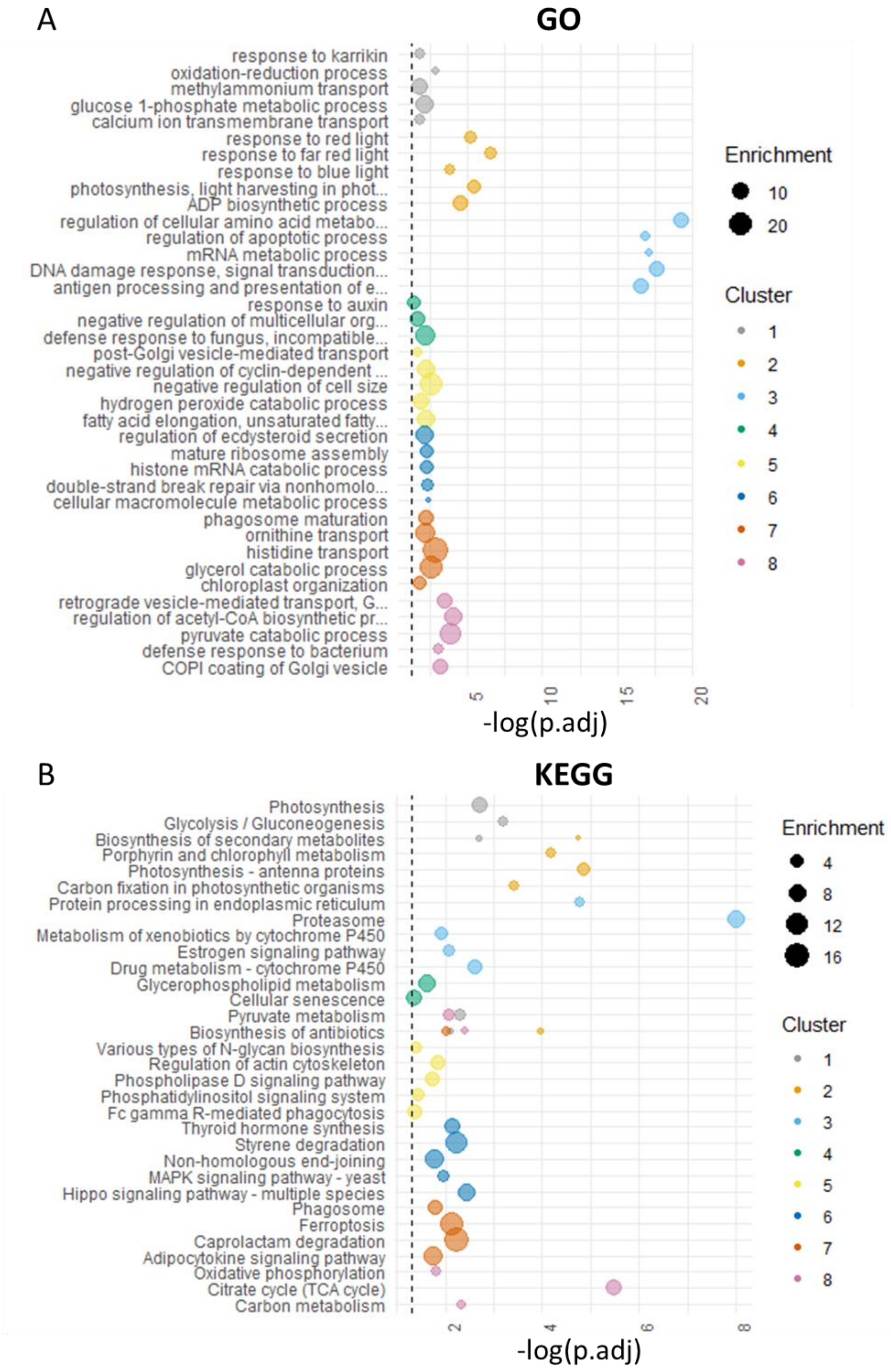


**Figure S3. GO and KEGG enrichment analysis.** The top 5 GO biological process (***A***) and KEGG (***B***) terms found to be significantly enriched (p.adj < 0.05) per cluster (in Fig. 1D) using a hypergeometric test are shown, over the –log10 adjusted *P*-value. Size represents enrichment, color represents the cluster. Dashed line represents the adjusted *P*-value threshold of 0.05.

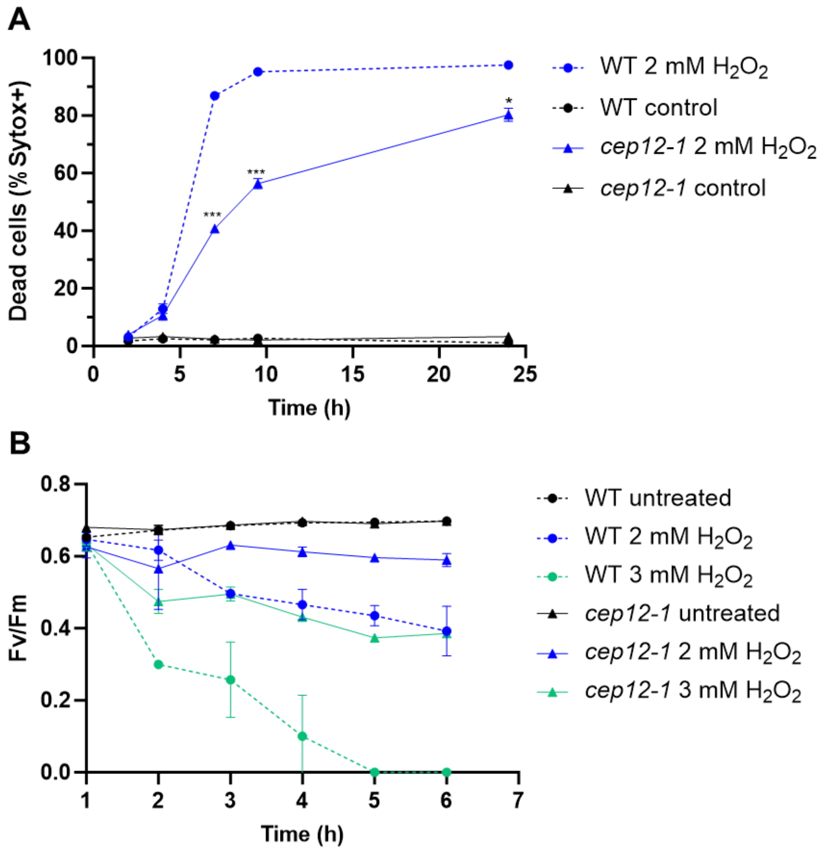


**Figure S4. *C. reinhardtii cep12-1* mutant is resilient to oxidative stress.** The mortality and maximal photosynthetic efficiency of *C. reinhardtii cep12-1* mutant (solid lines, triangles) and WT (dashed, circles) cells in response to $H_2O_2$. (***A***) Percentage of dead cells over time after $H_2O_2$ treatment (2 mM, blue) and in untreated control (black) as measured by positive Sytox Green staining using flow cytometry. (***B***) PSII maximal efficiency (Fv/Fm) over time after 2 mM (blue) and 3 mM (cyan) $H_2O_2$ treatments and in control untreated cells (black). (***A-B***) Values represent the mean ± sd, N=3. Statistical significance was calculated by Two-way repeated measures ANOVA as compared to WT. Asterisks represent *P*-values: * < 0.05, *** < 0.001.

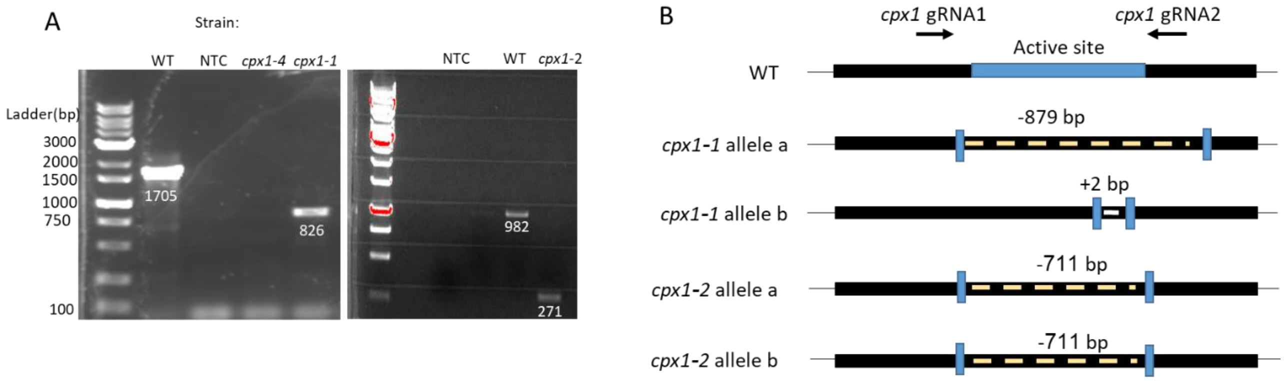


**Figure S5. CRISPR-Cas9 knockout of *CPX1* in *P. tricornutum*.** ***(A)*** Validation of deletions within the *CPX1* gene in *P. tricornutum cpx1* mutants by gel electrophoresis of PCR products using CPX1 primers. ***(B)*** Schematic representation of deletion in the DNA sequence of both alleles of the *CPX1* gene in the mutants and the WT. The blue area in the WT section represents the active site. Black arrows represent location of the two gRNAs. Deletion is marked in dashed yellow lines in the DNA. Clone *cpx1-1* has a deletion of 879 bp in allele a and an insertion of 2 bp in allele b, and clone *cpx1-2* has a 711 bp deletion in both alleles (see also Fig. S6).

```
wt        CGGGACAGCTTTTCTTCGATCATCATACCGGGATTACGAAATCAGCCAGACCGTTTCACG
cpx1-1a   CGGGACAG----------------------------------------------------

wt        atggccattcgaagcttcgttaatcttcaggttgtattcgtccttcttgctcttctatcg 60
cpx1-1a   ------------------------------------------------------------
cpx1-1b   atggccattcgaagcttcgttaatcttcaggttgtattcgtccttcttgctcttctatcg
cpx1-2    atggccattcgaagcttcgttaatcttcaggttgtattcgtccttcttgctcttctatcg
wt        M  A  I  R  S  F  V  N  L  Q  V  V  F  V  L  L  A  L  L  S   20
cpx1-1b   M  A  I  R  S  F  V  N  L  Q  V  V  F  V  L  L  A  L  L  S
cpx1-2    M  A  I  R  S  F  V  N  L  Q  V  V  F  V  L  L  A  L  L  S

wt        tgctcgttggattgggctgatgctcaagacgatgagcgcatcatgcccggtcacactcga 120
cpx1-1a   ------------------------------------------------------------
cpx1-1b   tgctcgttggattgggctgatgctcaagacgatgagcgcatcatgcccggtcacactAga
cpx1-2    tgctcgttggattgggctgatgctcaagacgatgagcgcatcatgccc------------
wt        C  S  L  D  W  A  D  A  Q  D  D  E  R  I  M  P  G  H  T  R   40
cpx1-1b   C  S  L  D  W  A  D  A  Q  D  D  E  R  I  M  P  G  H  T  R
cpx1-2    C  S  L  D  W  A  D  A  Q  D  D  E  R  I  M  P  -  -  -  -

wt        ctagaacacgtcgtcagcccactaccttacacttacctttcgaatgaggaactcccgatg 180
cpx1-1a   ------------------------------------------------------------
cpx1-1b   ctagaacacgtcgtcagcccactaccttacacttacctttcgaatgaggaactcccgatg
cpx1-2    ------------------------------------------------------------
wt        L  E  H  V  V  S  P  L  P  Y  T  Y  L  S  N  E  E  L  P  M   60
cpx1-1b   L  E  H  V  V  S  P  L  P  Y  T  Y  L  S  N  E  E  L  P  M
cpx1-2    -  -  -  -  -  -  -  -  -  -  -  -  -  -  -  -  -  -  -  -

wt        gccttttcatggggcaatgtcaatgggcgttcgtatttgaccaagtcattgaatcaacac 240
cpx1-1a   ------------------------------------------------------------
cpx1-1b   gccttttcatggggcaatgtcaatgggcgttcgtatttgaccaagtcattgaatcaacac
cpx1-2    ------------------------------------------------------------
wt        A  F  S  W  G  N  V  N  G  R  S  Y  L  T  K  S  L  N  Q  H   80
cpx1-1b   A  F  S  W  G  N  V  N  G  R  S  Y  L  T  K  S  L  N  Q  H
cpx1-2    -  -  -  -  -  -  -  -  -  -  -  -  -  -  -  -  -  -  -  -
```

```
wt         attcctcaatattgcggcaggtaggttaattattgaaaggtttctttcagtagcaatacc 300
cpx1-1a    ------------------------------------------------------------
cpx1-1b    attcctcaatattgcggcaggtaggttaattattgaaaggtttctttcagtagcaatacc
cpx1-2     ------------------------------------------------------------
wt         I  P  Q  Y  C  G  S                                          87
cpx1-1b    I  P  Q  Y  C  G  S
cpx1-2     -  -  -  -  -  -  -

wt         aaattacaaaattgattccaaaataatacattatccacctttttctcgctacagctgctg 360
cpx1-1a    ------------------------------------------------------------
cpx1-1b    aaattacaaaattgattccaaaataatacattatccacctttttctcgctacagctgctg
cpx1-2     ------------------------------------------------------------
wt                                                               C  W   89
cpx1-1b                                                          C  W
cpx1-2                                                           -  -

wt         ggcgcatgccgctttatcggttctcggggatcggataatgattgcccaaagccaagaaga 420
cpx1-1a    ------------------------------------------------------------
cpx1-1b    ggcgcatgccgctttatcggttctcggggatcggataatgattgcccaaagccaagaaga
cpx1-2     ------------------------------------------------------------
wt          A  H  A  A  L  S  V  L  G  D  R  I  M  I  A  Q  S  Q  E  E  109
cpx1-1b     A  H  A  A  L  S  V  L  G  D  R  I  M  I  A  Q  S  Q  E  E
cpx1-2      -  -  -  -  -  -  -  -  -  -  -  -  -  -  -  -  -  -  -  -

wt         ggactctagcattctcgatgagtttaatctttcggttcagtttctattaaattgcgctgg 480
cpx1-1a    ------------------------------------------------------------
cpx1-1b    ggactctagcattctcgatgagtttaatctttcggttcagtttctattaaattgcgctgg
cpx1-2     ------------------------------------------------------------
wt          D  S  S  I  L  D  E  F  N  L  S  V  Q  F  L  L  N  C  A  G  129
cpx1-1b     D  S  S  I  L  D  E  F  N  L  S  V  Q  F  L  L  N  C  A  G
cpx1-2      -  -  -  -  -  -  -  -  -  -  -  -  -  -  -  -  -  -  -  -

wt         cgaatatgccggatcttgttacggaggatcgacaacaggagtgttcgacttcatccaaga 540
cpx1-1a    ------------------------------------------------------------
cpx1-1b    cgaatatgccggatcttgttacggaggatcgacaacaggagtgttcgacttcatccaaga
cpx1-2     ------------------------------------------------------------
wt          E  Y  A  G  S  C  Y  G  G  S  T  T  G  V  F  D  F  I  Q  D  149
cpx1-1b     E  Y  A  G  S  C  Y  G  G  S  T  T  G  V  F  D  F  I  Q  D
cpx1-2      -  -  -  -  -  -  -  -  -  -  -  -  -  -  -  -  -  -  -  -

wt         catggggtacattccctatgagacttgtcagccgtaccttgcctgctccgatgattctga 600
cpx1-1a    ------------------------------------------------------------
cpx1-1b    catggggtacattccctatgagacttgtcagccgtaccttgcctgctccgatgattctga
cpx1-2     ------------------------------------------------------------
wt          M  G  Y  I  P  Y  E  T  C  Q  P  Y  L  A  C  S  D  D  S  D  169
cpx1-1b     M  G  Y  I  P  Y  E  T  C  Q  P  Y  L  A  C  S  D  D  S  D
cpx1-2      -  -  -  -  -  -  -  -  -  -  -  -  -  -  -  -  -  -  -  -

wt         cgaaggtatttgttcttttgtaaataccacgtgctcacccgaagcaatttgccgtacatg 660
cpx1-1a    ------------------------------------------------------------
cpx1-1b    cgaaggtatttgttcttttgtaaataccacgtgctcacccgaagcaatttgccgtacatg
cpx1-2     ------------------------------------------------------------
wt          E  G  I  C  S  F  V  N  T  T  C  S  P  E  A  I  C  R  T  C  189
cpx1-1b     E  G  I  C  S  F  V  N  T  T  C  S  P  E  A  I  C  R  T  C
cpx1-2      -  -  -  -  -  -  -  -  -  -  -  -  -  -  -  -  -  -  -  -

wt         ctctcccgacggcatttgccaagctgttaccactttcccgaacgccactgtcgctgagta 720
cpx1-1a    ------------------------------------------------------------
cpx1-1b    ctctcccgacggcatttgccaagctgttaccactttcccgaacgccactgtcgctgagta
cpx1-2     ------------------------------------------------------------
wt          S  P  D  G  I  C  Q  A  V  T  T  F  P  N  A  T  V  A  E  Y  209
cpx1-1b     S  P  D  G  I  C  Q  A  V  T  T  F  P  N  A  T  V  A  E  Y
cpx1-2      -  -  -  -  -  -  -  -  -  -  -  -  -  -  -  -  -  -  -  -

wt         tggccggtaccgttatgaactgtttgctactatggcggaaatctatctccgtggtccagt 780
cpx1-1a    ------------------------------------------------------------
cpx1-1b    tggccggtaccgttatgaactgtttgctactatggcggaaatctatctccgtggtccagt
cpx1-2     ------------------------------------------------------------
wt          G  R  Y  R  Y  E  L  F  A  T  M  A  E  I  Y  L  R  G  P  V  229
cpx1-1b     G  R  Y  R  Y  E  L  F  A  T  M  A  E  I  Y  L  R  G  P  V
```

```
cpx1-2    -  -  -  -  -  -  -  -  -  -  -  -  -  -  -  -  -  -  -  -

wt        tactgcatccattgatgccggaccaattcacaaata--ccctggtggtgtcttgtgggataa 840
cpx1-1a   ---------------------------------------------------tcttgtgggataa
cpx1-1b   tactgcatccattgatgccggaccaattcacaaataTAccctggtggtgtcttgtgggataa
cpx1-2    -------------------------------------------tggtggtgtcttgtgggataa
wt         T  A  S  I  D  A  G  P  I  H  K  Y     P  G  G  V  L  W  D  N  249
cpx1-1b    T  A  S  I  D  A  G  P  I  H  K  Y  T  L  V  V  S  C  G  I  I
cpx1-2     -  -  -  -  -  -  -  -  -  -  -  -  -   W  W  C  L  V  G  *

wt        tcccaaatatcattccgacaagacaaaccacgctgttagcattgttggctggggctacga 900
cpx1-1a   tcccaaatatcattccgacaagacaaaccacgctgttagcattgttggctggggctacga
cpx1-1b   tcccaaatatcattccgacaagacaaaccacgctgttagcattgttggctggggctacga
cpx1-2    tcccaaatatcattccgacaagacaaaccacgctgttagcattgttggctggggctacga
wt         P  K  Y  H  S  D  K  T  N  H  A  V  S  I  V  G  W  G  Y  D  269
cpx1-1b     P  N  I  I  P  T  R  Q  T  T  L  L  A  L  L  A  G  A  T  I

wt        ttacgatgaagagaagcagtactggatcgttcgaaactcttggggacaatattggggtga 960
cpx1-1a   ttacgatgaagagaagcagtactggatcgttcgaaactcttggggacaatattggggtga
cpx1-1b   ttacgatgaagagaagcagtactggatcgttcgaaactcttggggacaatattggggtga
cpx1-2    ttacgatgaagagaagcagtactggatcgttcgaaactcttggggacaatattggggtga
wt         Y  D  E  E  K  Q  Y  W  I  V  R  N  S  W  G  Q  Y  W  G  E  289
cpx1-1b     T  M  K  R  S  S  T  G  S  F  E  T  L  G  D  N  I  G  V  K

wt        aatgggattctttcgtatcgagcttggtaagaatctgctcaagatagaatccaacattgc 1020
cpx1-1a   aatgggattctttcgtatcgagcttggtaagaatctgctcaagatagaatccaacattgc
cpx1-1b   aatgggattctttcgtatcgagcttggtaagaatctgctcaagatagaatccaacattgc
cpx1-2    aatgggattctttcgtatcgagcttggtaagaatctgctcaagatagaatccaacattgc
wt         M  G  F  F  R  I  E  L  G  K  N  L  L  K  I  E  S  N  I  A  309
cpx1-1b     W  D  S  F  V  S  S  L  V  R  I  C  S  R  *

wt        ttgggcaaaccccgggaccttctctgtctttgatcctgagtgtacagatggagattgccg 1080
cpx1-1a   ttgggcaaaccccgggaccttctctgtctttgatcctgagtgtacagatggagattgccg
cpx1-1b   ttgggcaaaccccgggaccttctctgtctttgatcctgagtgtacagatggagattgccg
cpx1-2    ttgggcaaaccccgggaccttctctgtctttgatcctgagtgtacagatggagattgccg
wt         W  A  N  P  G  T  F  S  V  F  D  P  E  C  T  D  G  D  C  R  329

wt        cctgcgttcgatgacttacactgatccgtctcaagatatccagttgttagagcaaagact 1140
cpx1-1a   cctgcgttcgatgacttacactgatccgtctcaagatatccagttgttagagcaaagact
cpx1-1b   cctgcgttcgatgacttacactgatccgtctcaagatatccagttgttagagcaaagact
cpx1-2    cctgcgttcgatgacttacactgatccgtctcaagatatccagttgttagagcaaagact
wt         L  R  S  M  T  Y  T  D  P  S  Q  D  I  Q  L  L  E  Q  R  L  349

wt        tcgtatcggagtgcaattctga 1162
cpx1-1a   tcgtatcggagtgcaattctga
cpx1-1b   tcgtatcggagtgcaattctga
cpx1-2    tcgtatcggagtgcaattctga
wt         R  I  G  V  Q  F  *   355
```

**Figure S6. Sequences of *CPX1* in *P. tricornutum* mutants***.* DNA and protein sequences of WT *P. tricornutum CPX1* and the two *cpx1* mutants. The one letter code for the amino acid is directly below the first base of the codon. Blue letters represent untranslated regions (upper case – upstream untranslated region, lower case – introns), black lowercase letters are exons. Cyan background represents the guide sequence. Catalytic site residues are in pink with grey background. Difference between the WT and the mutants in both DNA and protein are in red. In *cpx1-1* allele b, the insertion of two base pairs (marked in red) caused a frame shift leading to a premature stop codon. In *cpx1-2*, both alleles had the same 711 bp deletion leading to six novel amino acids after the 36 amino acids in common with the wild type, and 319 amino acids are missing due to the deletion. Catalytic site residues H259 and N281 do not exist in both *cpx1-1* and *cpx1-2*.

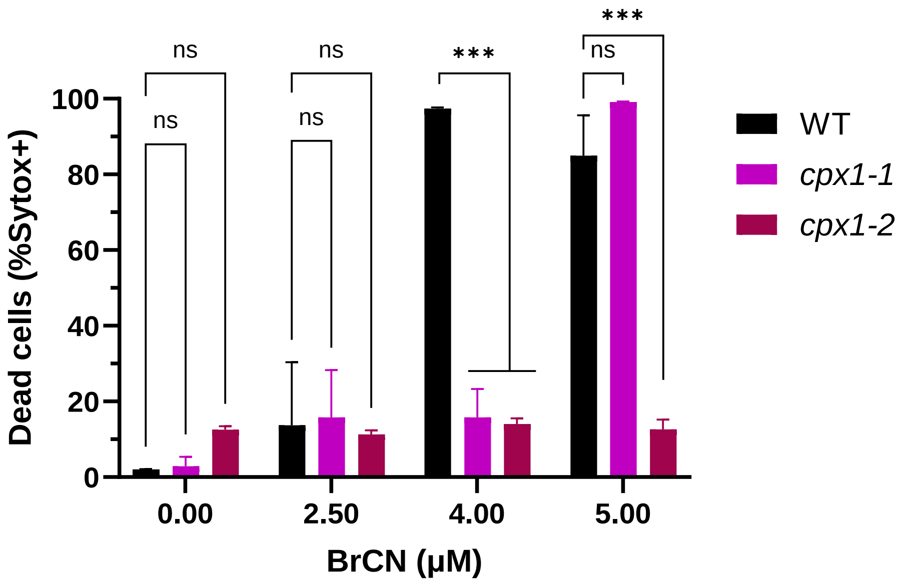


**Figure S7. *P. tricornutum cpx1* mutants are resilient to BrCN.** *P. tricornutum* WT (black) and *cpx1* mutants (purple and red) were treated with different doses of BrCN, and cell death was measured 24 h later using Sytox green staining and flow cytometry. Values represent the mean ± sd of biological triplicates. Statistical significance was calculated using two-way ANOVA as compared to WT cells. Asterisks represent *P*-values < 0.001. ns = non-significant.

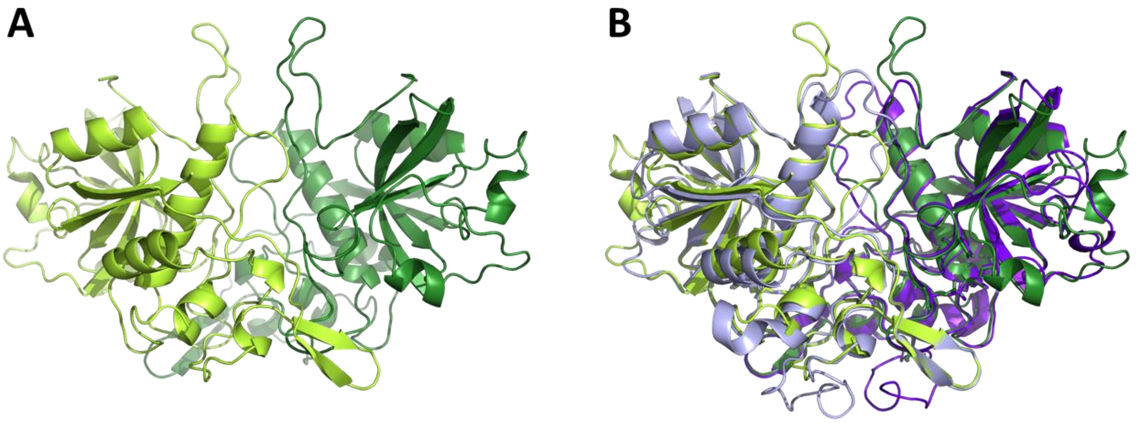


**Figure S8. Structure models of *C. reinhardtii* CEP12, *P. tricornutum* CPX1 and *Homo sapiens* CTSZ.** Overlays of AlphaFold2 predicted model of *C. reinhardtii* CEP12 (green) with the experimental structure of human CTSZ (PDB ID 1EF7, grey, **A**) and with the predicted model of *P. tricornutum* CPX1 (purple, **B**) dimers. The monomers in each homodimer are shown in light and dark shades.

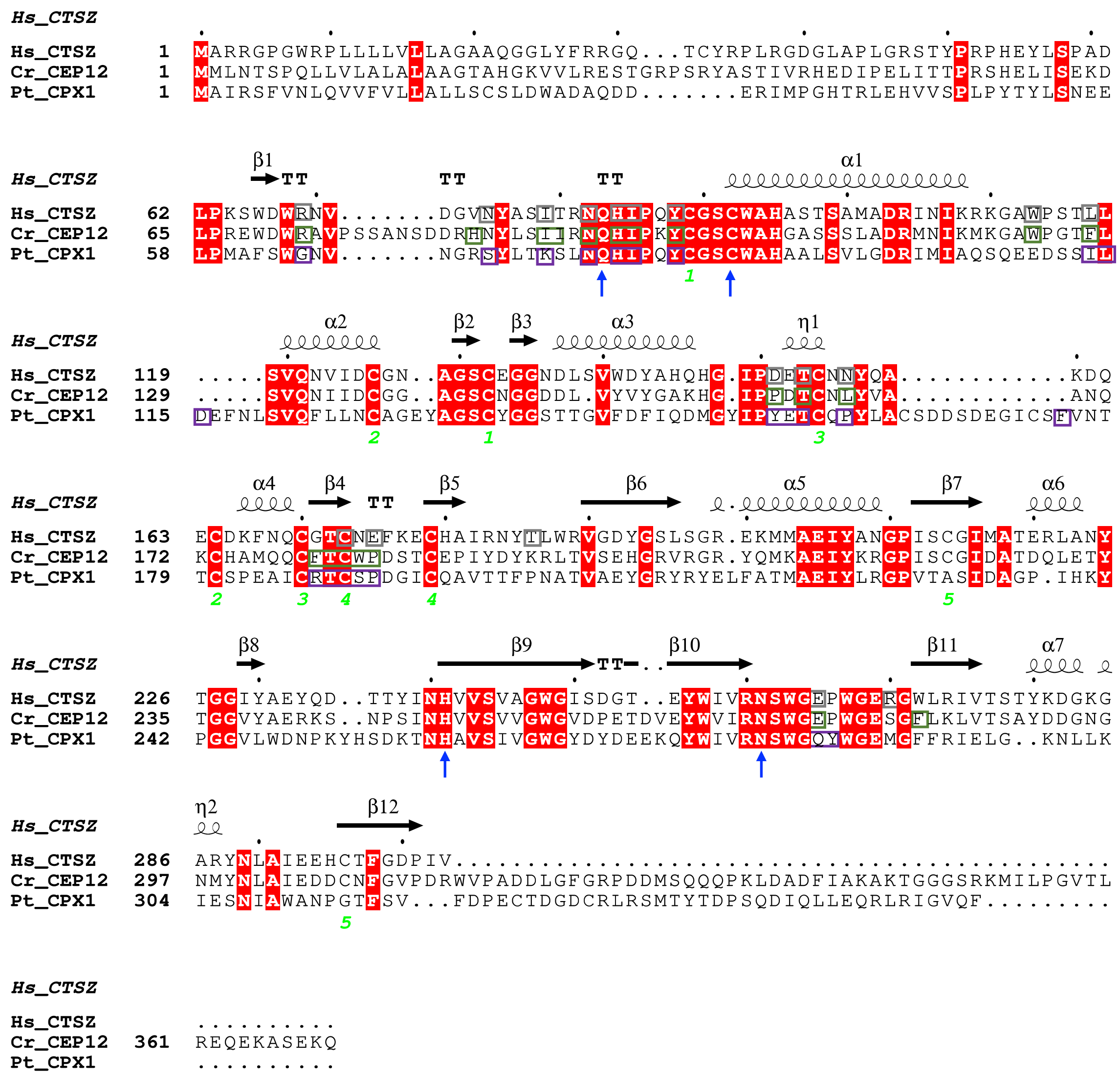


**Figure S9. Sequence alignment of CPX.** Protein sequence alignment of *H. sapiens* CTSZ (Hs_CTSZ, PDB ID 1EF7), *C. reinhardtii* CEP12 (Cr_CEP12) and *P. tricornutum* CPX1 (Pt_CPX1). Secondary structure elements of Hs_CTSZ are labelled above the alignments: α-helices and $3_{10}$-helices (shown with the symbol "η") are indicated by coils, and β-strands by arrows. The residues conserved in all variants are in red and green numbers indicate the S-S bond between pairs of Cys residues. Active site residues are shown in blue arrows and residues involve in dimer interface are shown in boxes (grey – Hs_CTSZ; green – Cr_CEP12; purple – Pt_CPX1). The figure was created using ESPript [2].

**A**

```
>contig_520908_188_1195_-
Length=1008

 Score = 344 bits (883),  Expect = 1e-118, Method: Compositional matrix adjust.
 Identities = 173/327 (53%), Positives = 220/327 (67%), Gaps = 22/327 (7%)
 Frame = +1

Pt_CPX1  37   GHTRLEHVVSPLPYTYLSNEELPMAFSWGNVNGRSYLTKSLNQHIPQYCGSCWAHAALSV  96
              G  R +H    PLP+TY+    +LP +F+W +V+G SYLTKSLNQHIPQYCGSCWAH ALS
PaCPX1   1    GKIRHQH---PLPHTYIEAGDLPQSFNWADVDGVSYLTKSLNQHIPQYCGSCWAHGALSS  171

Pt_CPX1  97   LGDRIMIAQSQEEDSSILDEFNLSVQFLLNCAGEYAGSCYGGSTTGVFDFIQDMGYIPYE  156
              LGDRI IA+  + D      + NLS+QF+LNC  E AGSC+GG  T  ++FIQ  GY+PY+
PaCPX1   172  LGDRIKIARKAKGD-----DINLSIQFILNCGTESAGSCHGGYHTSTYEFIQTTGYVPYD  336

Pt_CPX1  157  TCQPYLACSDDSDEGICSFVNTTCSPEAICRTCSP----DGICQAVTTFPNATVAEYGRY  212
              TC PYLACS +S EG C  ++TTC+ E  C+TC        G C A+   FPNATVAEYG
PaCPX1   337  TCTPYLACSSESTEGFCPQIDTTCTAENTCKTCDTFGGMGGKCVALDYFPNATVAEYGLI  516

Pt_CPX1  213  RYE-------LFATMAEIYLRGPVTASIDAGPIHKYPGGVLWDNPKYHSDKTNHAVSIVG  265
               Y+       +   MAEIY RGPV A+I+A PI KY GG+   +    +S+ TNH V+IVG
PaCPX1   517  DYDADNKEDTVHKIMAEIYSRGPVAATINAEPIVKYTGGIFAETG--YSEDTNHIVAIVG  690

Pt_CPX1  266  WGYDYDEEKQYWIVRNSWGQYWGEMGFFRIELGKNLLKIESNIAWANPGTFSVFDPEC-T  324
              WG D + + Q+WIVRNSWGQYWGEMG+ R+E+GKNLL IE  IAWA PG F+  +  C
PaCPX1   691  WGTDKETKAQHWIVRNSWGQYWGEMGYMRLEMGKNLLGIEGEIAWAVPGEFTTTNYACYE  870

Pt_CPX1  325  DGDCRLRSMTYTDPSQDIQLLEQRLRI  351
              DG   + +  Y DPS D+  ++ RL +
PaCPX1   871  DGSNCVHTEEYVDPSTDVVAMKHRLFV  951
```

**B**

```
>contig_389670_1124_2455_-
Length=1332

 Score = 332 bits (851),  Expect = 3e-112, Method: Compositional matrix adjust.
 Identities = 168/330 (51%), Positives = 215/330 (65%), Gaps = 20/330 (6%)
 Frame = +1

Pt_CPX1  34    IMPGHTRLEHVVSPLPYTYLSNEELPMAFSWGNVNGRSYLTKSLNQHIPQYCGSCWAHAA  93
               ++ GHT+ E+  SPLP+TY+  E+LP    W NV+G SY T SLNQHIPQYCGSCWAH A
PaCPX2   310   VLEGHTQRENYHSPLPHTYIKEEDLPDNLDWRNVDGESYTTHSLNQHIPQYCGSCWAHGA  489

Pt_CPX1  94    LSVLGDRIMIAQSQEEDSSILDEFNLSVQFLLNCAGEYAGSCYGGSTTGVFDFIQDMGYI  153
               LS LGDRI IA++        DE NLS+Q++LNCA   AGSC+GGS TGVF+FIQ  G +
PaCPX2   490   LSSLGDRIKIARN-----GTGDEINLSIQYILNCATATAGSCHGGSHTGVFEFIQKSGLV  654

Pt_CPX1  154   PYETCQPYLACSDDSDEGICSFVNTTCSPEAICRTCSP----DGICQAVTTFPNATVAEY  209
               PY+TCQPYLACS +S EG C  V+TTCS     C+TC        G C  +    PNAT+ EY
PaCPX2   655   PYDTCQPYLACSSESTEGFCPHVDTTCSKLNTCKTCDTFGGMGGQCTEIDVMPNATIGEY  834

Pt_CPX1  210   GRYRYE------LFATMAEIYLRGPVTASIDAGPIHKYPGGVLWDNPKYHSDKTNHAVSI  263
               G Y +        +    +EIY RGPV    ++A PI +Y GG + D+ K+      NH VSI
PaCPX2   835   GTYSFLSDPSGIVHQIQSEIYARGPVATGVNAEPIVEYTGGRV-DDTKFWHMMVNHIVSI  1011

Pt_CPX1  264   VGWGYDYDEEKQYWIVRNSWGQYWGEMGFFRIELGKNLLKIESNIAWANPGTFSVFDPEC  323
               VGW   D +     YWIVRNSWGQYWGEMG+FRI   G N L IE ++AWA PG F+V +  C
PaCPX2   1012  VGWETDEETGNVYWIVRNSWGQYWGEMGYFRILAGHNSLGIEMDVAWATPGQFTVHNFPC  1191

Pt_CPX1  324   TD--GDCRLRSMT--YTDPSQDIQLLEQRL  349
                +   +C    + T   Y DPS+++++++ RL
PaCPX2   1192  YENGANCNGGATTQFYEDPSKNVEMVQARL  1281
```

**Figure S10. Sequence alignment of *P. tricornutum CPX1* and *P. australis* contigs.** Protein sequence alignment of *P. tricornutum* CPX1 and *P. australis* contigs PaCPX1 (contig_520908_188_1195_-; **A**) and PaCPX2 (contig_389670_1124_2455_-; **B**) from metatrascriptomics data of a *P. australis* dominated bloom[3].

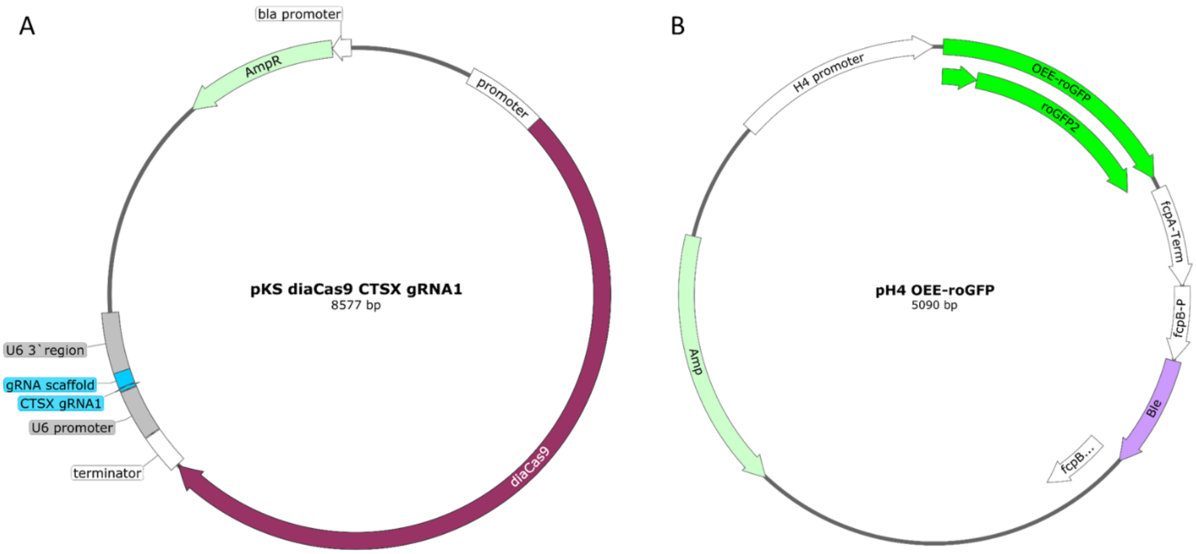


**Figure S11. Plasmids used for generating *P. tricornutum* CPX1 mutants using CRISPR-Cas9.** To generate *P. tricornutum* mutants with large deletions in the CPX1 gene, cells were co-transformed with 3 plasmids using biolistic bombardment: 2 plasmids containing diaCas9 and CPX1 gRNA targeting different regions of the CPX1 gene (gRNA1 and gRNA2 in each plasmid, as in plasmid pKS diaCas9 CTSX gRNA1 shown in ***A***), and a plasmid containing the zeocin resistance cassette and chloroplast targeted roGFP (pH4 OEE-roGFP, ***B***). We did not detect chl-roGFP fluorescent signal in *P. tricornutum cpx1-1* and *cpx1-2* clones, potentially due to low expression or silencing of the chl-roGFP construct or partial integration of the plasmid depicted in B.

### References – supplementary information


1. Schwarzländer, M. *et al.* Confocal imaging of glutathione redox potential in living plant cells. *J. Microsc.* **231**, 299–316 (2008).

2. Robert, X. & Gouet, P. Deciphering key features in protein structures with the new ENDscript server. *Nucleic Acids Res.* **42**, W320–W324 (2014).

3. Brunson, J. K. *et al.* Molecular Forecasting of Domoic Acid during a Pervasive Toxic Diatom Bloom. *bioRxiv* (2023).
